# Oncogenic KRAS cells use Wnt signalling and cell dormancy to override homeostatic cell elimination mechanisms in adult pancreas

**DOI:** 10.1101/2024.02.13.579930

**Authors:** Beatriz Salvador-Barbero, Markella Alatsatianos, Jennifer P. Morton, Owen J. Sansom, Catherine Hogan

**Affiliations:** European Cancer Stem Cell Research Institute, School of Biosciences, Cardiff University, Cardiff CF24 4HQ, UK; CRUK Scotland Institute, Glasgow, G61 1BD, UK; School of Cancer Sciences, University of Glasgow, Glasgow, G61 1QH, UK

**Keywords:** Epithelial homeostasis, cell extrusion, cell competition, oncogenic RAS, mutant Tp53, pancreas, Wnt5a, cell dormancy, pancreatic cancer

## Abstract

Epithelial tissues use homeostatic defence mechanisms to actively expel aberrant or genetically mutant cells and prevent disease. When present in healthy tissues in low numbers, we show that cells expressing cancer-causing mutations (KrasG12D, p53R172H) compete with normal cells for survival and are often eliminated. Thus, tumour initiation must require mechanisms whereby mutant cells override tissue defence mechanisms to remain in a tissue; however, the biology of these initial events is poorly understood. Here, we use an *in vivo* model of sporadic tumorigenesis in the adult pancreas to show that a population of KrasG12D- or p53R172H-expressing cells are never eliminated from the epithelium. Using RNA sequencing of non-eliminated populations and quantitative fluorescence imaging, we show that β-catenin-independent Wnt5a signalling, and cell dormancy are key features of surviving KrasG12D cells *in vivo*. We demonstrate that Wnt5a (and not Wnt3a) inhibits apical extrusion of RasV12 cells *in vitro* by promoting stable E-cadherin-based cell-cell adhesions at RasV12-normal cell-cell boundaries. Inhibition of Wnt5a signalling restores E-cadherin dynamics at normal-mutant boundaries and apical extrusion *in vitro*. RasV12 cells arrested in the cell cycle are not extruded and this is rescued when Wnt signalling is inhibited. In the pancreas, Wnt signalling, E-cadherin and β-catenin are increased at cell-cell contacts between non-eliminated KrasG12D cells and normal neighbours. Importantly, we demonstrate that active Wnt signalling is a general mechanism required to promote KrasG12D and p53R172H cell survival *in vivo*. Treatment with porcupine inhibitor rescues pancreas tissue defence by switching mutant cell retention to cell expulsion. Our results suggest that RAS mutant cells activate Wnt and a dormant cell state to avoid cell expulsion and to survive in the adult pancreas.

## Introduction

Epithelial tissues are continuously exposed to mutational insults, and as a result, genetically mutant cells often arise in a tissue. Indeed, most human cancers start sporadically from epithelial cells carrying genetic mutations that activate oncogenes or inactivate tumour suppressor function. Remarkably, epithelial tissues protect against tumorigenesis by actively removing aberrant (*1, 2*) and genetically mutant cells (*3–9*). In general, genetically different cells compete for space and survival in tissues, resulting in the elimination of ‘less fit’ mutant cells via apoptosis, extrusion and/or cell differentiation (*10*). Cell competition outcomes are dependent on the genetic status of a cell relative to its local neighbourhood (*10, 11*) and extrinsic environmental factors (*12–14*) and therefore under certain conditions, competition is tipped in favour of mutant cells. This is better understood in rapidly proliferating tissues that are actively replenished via defined stem cell compartments; tumour risk increases when stem cells acquire genetic mutations that confer a competitive advantage over wild type counterparts, allowing the mutant stem cells to completely occupy the niche (*15–18*). In contrast, our understanding of how cell competition outcomes shape tumorigenesis in slow proliferating adult tissues that lack a *bona fide* stem cell compartment remains poorly detailed.

Using mouse pancreas as a model system of a slow proliferating adult tissue, we recently demonstrated that cells expressing oncogenic *Kras* (*Kras*^G12D^) are outcompeted by healthy neighbours and are eliminated from exocrine and endocrine epithelial compartments (*4*). We have previously described how interactions with normal cells trigger robust evolutionary conserved cell biology phenotypes in *Ras* transformed cells, resulting in the elimination of the mutant cells via cell segregation and cell extrusion (*4, 5, 9, 19*). Removal of KrasG12D cells from adult pancreas tissues requires local remodelling of E-cadherin-based cell-cell adhesions at normal-mutant boundaries and dynamic changes in mutant and normal cell volume (*4*). Importantly, we demonstrated that abrogation of KrasG12D cell elimination in the pancreas significantly increased the appearance of preneoplastic lesions (*4*), suggesting cell competition and the subsequent expulsion of mutant cells are disease preventative.

Activating mutations in oncogenic *RAS* are the principal driver gene event in human pancreatic ductal adenocarcinoma (PDAC; (*20*)), the most common type of human pancreatic cancer. *KRAS* mutations are detected in >90% of human tumours and *in vivo* mouse studies show that KRAS signalling is essential for disease to progress through all stages (*21*). Missense mutations in *TP53* are the second most common mutation detected in ∼70% of human PDAC (*22*) and required for metastasis (*23*). How genetically mutant cells survive and grow in the competitive environment of the adult pancreas remains unclear. Here, we sought to understand the mechanisms underpinning how mutant cells avoid cell expulsion and survive in the adult pancreas. Using *in vivo* models of sporadic tumorigenesis, we confirm that adult pancreas tissues actively eliminate cells expressing either oncogenic KrasG12D or p53R172H mutations; however, some mutant cells are never eliminated, suggesting these cells have a survival advantage. We report that cell competition is completely abrogated when tissues contain ‘double mutant’ cells, i.e., cells expressing both KrasG12D and p53R172H mutations. Using bulk RNA sequencing, we identify gene signatures of ‘never-eliminated’ mutant cells. We demonstrate a functional role of Wnt signalling as an important general regulator of mutant cell fate and cell dormancy as a second mechanism, whereby KrasG12D cells avoid cell elimination.

## Results

### Cells expressing p53R172H mutations are outcompeted and eliminated from adult pancreas compartments. Cells co-expressing p53172H and KrasG12D mutations are never eliminated

To model sporadic pancreatic cancer, we used the pancreas-specific genetically engineered mouse models: KC: *Pdx1-Cre^ERT^; LSL-Kras^G12D/+^; Rosa26^LSL-tdRFP^*; PC: *Pdx1-Cre^ERT^; LSL-Trp53^R172H/+^; Rosa26^LSL-tdRFP^*; and KPC: *Pdx1-Cre^ERT^; LSL-Kras^G12D/+^; Trp53^R172H/+^; Rosa26^LSL-tdRFP^*. Our experimental control was *Pdx1-Cre^ERT^; Rosa26^LSL-tdRFP^* mice (Fig. 1A left). Adult mice were treated with low-dose tamoxifen to induce transgene expression (together with RFP reporter) in a mosaic manner in all pancreatic epithelial compartments (*4*). To monitor genetically mutant cell fate, we measured RFP levels in tissues harvested at 7-, 35- and 70-days post induction (p.i.) of Cre recombinase (Fig. 1A-1C). Consistent with our previously published data (*4*), we found that low dose tamoxifen induced stochastic RFP labelling in ∼20-25% of the tissue (Fig. 1C); levels of RFP fluorescence were comparable between all genotypes at 7 days (Fig. 1B, top panels; 1C; (*4*)) and significantly decreased over time in KrasG12D tissues (KC; Fig. 1B, lower panel; 1C, *p*<0.0001 for both 35 and 70 days p.i.). RFP fluorescence also significantly decreased over time in tissues expressing p53R172H mutations (PC; Fig. 1B, lower panel; 1C, *p*<0.0001 and *p=*0.0173 at 35 and 70 days p.i. respectively). In contrast, we found no significant difference in the level of RFP in double mutant (KPC) tissues over time (Fig. 1B, lower panel; 1C, p=0.2062 and p=0.1504 at 35 and 70 days p.i. respectively). RFP+ ducts were significantly less frequent in KrasG12D (KC) and p53R172H (PC) tissues at the 7-day p.i. time point, compared to controls (Fig. 1D, *p=*0.0038 and *p*=0.0029 respectively), while the number of RFP+ ducts in double mutant (KPC) tissues was comparable to controls (Fig. 1D, *p*=0.2879). We also measured a significant decrease in the percentage of RFP+ cells per islet in p53R172H (PC) and KrasG12D (KC) tissues (Fig. 1E, *p*=0.0214 and *p*=0.0134 respectively), whereas the frequency of RFP+ cells per islet in double mutant (KPC) tissues was also comparable to wild type controls (Fig. 1E, *p*=0.5394). Thus, like KrasG12D-expressing cells (*4*), p53R172H single mutant cells are eliminated from all epithelial compartments. Cells expressing both KrasG12D and p53R172H mutations are never eliminated from endocrine/exocrine epithelial tissues.

**Figure 1:**
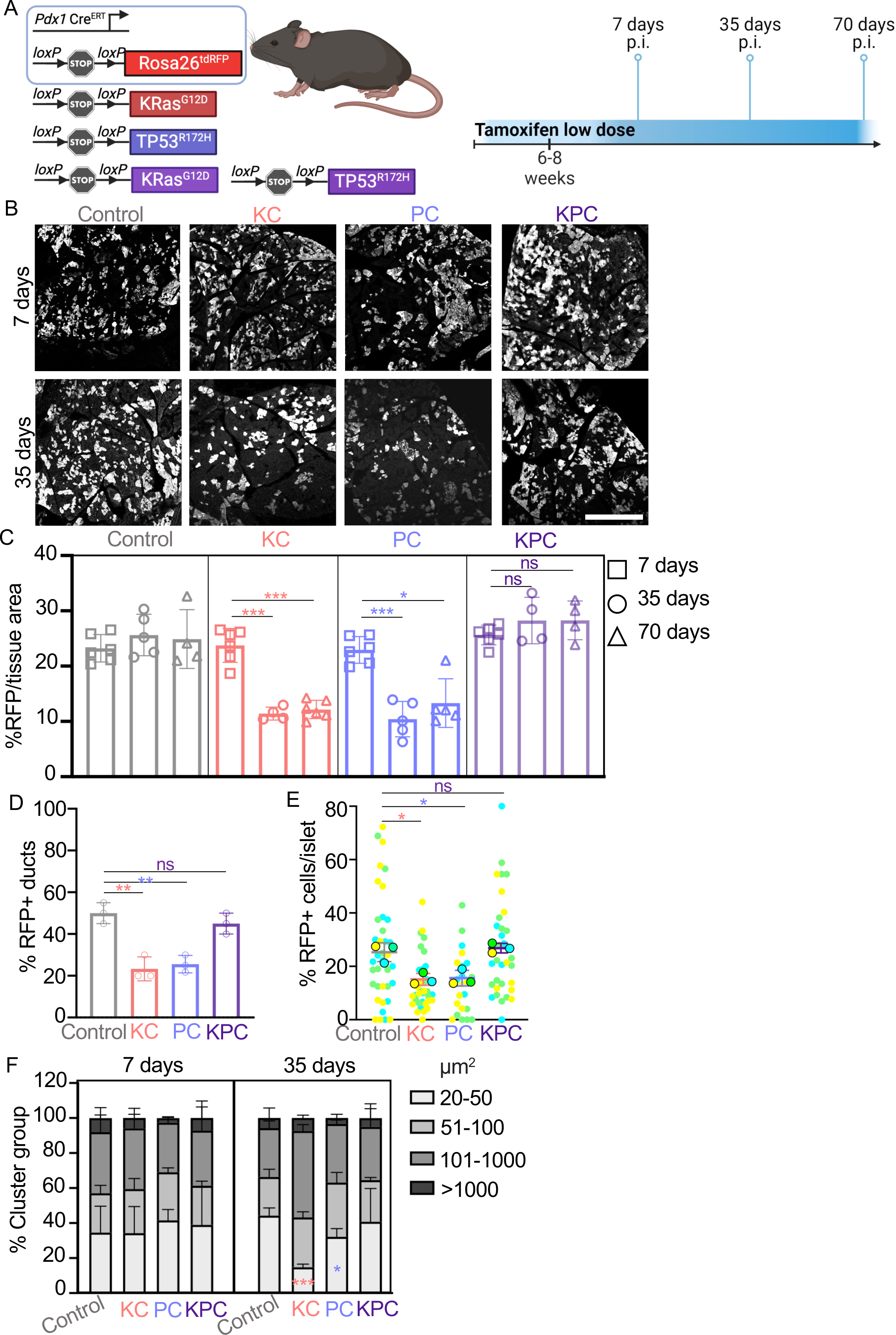
Cells expressing p53R172H are eliminated from adult pancreas *in vivo*. Co-expression of p53R172H mutations prevents KrasG12D cell elimination *in vivo*. **A** Schematic of mouse models (left), tamoxifen treatment and timepoints (right). 6–8-week-old *Pdx1-Cre^ERT^; Rosa26^LSL-tdRFP^* (Control), *Pdx1-Cre^ERT^; Rosa26^LSL-tdRFP^; LSL-Kras^G12D/+^*(KC), *Pdx1-Cre^ERT^; Rosa26^LSL-tdRFP^; LSL-TP53^R172H/+^*(PC), and *Pdx1-Cre^ERT^; Rosa26^LSL-tdRFP^; LSL-Kras^G12D/+^; LSL-TP53^R172H/+^* (KPC) mice were treated with one single low dose of tamoxifen. Pancreata were harvested at 7-, 35- or 70-days post tamoxifen induction. Illustration created with BioRender.com. **B** Representative images of endogenous RFP fluorescence showing the area occupied by RFP in Control, KrasG12D (KC), p53R172H (PC) or double mutant (KPC) pancreas tissue sections harvested at 7-days (top) and 35-days (bottom) p.i. Scale bar, 500µm. **C** Percentage of RFP fluorescence per total tissue area from Control, KrasG12D (KC), p53R172H (PC) or double mutant (KPC) tissues at 7-(squares), 35-(circles) and 70-days (triangles) p.i. Each data point represents average RFP fluorescence per mouse; control n=6 (7-days p.i.), n=5 (35-days p.i.), n=4 (70-days p.i.); KC n=6 (7- and 70-days p.i.), n=4 (35-days p.i.); PC n=6 (7-days p.i.), n=5 (35- and 70-days p.i.); KPC n=6 (7-days p.i.), n=4 (35- and 70-days p.i.). The data represent mean and +/- s.d. Student’s t test was used to analyse the data, \*\*\**p*<0.0001, \**p*=0.0173, ns>0.1504. **D** Percentage of RFP+ ducts at 7-days p.i. Ducts that carried at least one RFP+ cell were quantified as positive. Data points represent mean RFP+ percentage per mouse (n=3 per group), with at least 20 ducts counted per mouse. The data represent mean and +/- s.d. Student’s t test was used to analyse the data, \*\**p*<0.004, ns=0.2879. **E** Percentage of RFP+ cells per total number of cells in islets at 35 days p.i. SuperPlot shows all the quantified islets (smaller circles) and the mean for each mouse (n=3 samples; larger circles - blue, green, and yellow). The graph shows mean and +/- s.d. for the three mice. Student’s t test was used to analyse the data, \**p*<0.03, ns=0.5394. **F** Percentage of RFP positive clusters grouped according to cluster area (μm^2^). RFP positive cluster area was measured, and clusters were grouped according to their size in 4 groups at 7 and 35 days p.i.: 20-50 μm^2^, 51-100 μm^2^, 101-1000 μm^2^ and more than 1000 μm^2^. The graph shows the percentage of clusters in each group relative to the total number of clusters. At 35 days p.i. the percentage of 20-50 μm^2^ clusters significantly decreases in KC (*p*=0.0004) and PC (*p*=0.0316), no significant differences were observed in Control or KPC tissues (p=0.7740). Student’s t test was used to analyse the data. The data represent mean +/- s.d. for at least 700 clusters were quantified per mouse and 3 mice per genotype/timepoint.

We have previously shown that KrasG12D cells are competitively eliminated at cell boundaries with surrounding normal cells, resulting in the elimination of small clusters (*4, 24*). Similar to our previously published results (*4*), the percentage of small clusters (<50 micron^2^) significantly decreased in KrasG12D (KC; Fig. 1F, *p*=0.0004) and in p53R172H (PC; Fig. 1F, *p*=0.0316) tissues over time. We did not see significant differences in cluster distribution based on size in double mutant (KPC) tissues, compared to controls (Fig. 1F; *p*=0.7740, small clusters), suggesting double mutant (KPC) cells do not respond to cell-cell interactions with normal cells. Thus, expression of p53R172H mutations in low numbers of cells in adult pancreas tissues triggers cell competition *in vivo*, resulting in elimination of p53R172H cells. Co-expression of mutant p53R172H confers a survival advantage to KrasG12D cells, abrogating competition in the tissue and allowing KrasG12D cells to avoid cell elimination.

### Transcriptional profiles of non-eliminated mutant cells indicate activation of pro-survival signals

We consistently found that ∼10% of tissue area remains RFP-labelled in KrasG12D (KC) and in p53R172H (PC) tissues post the 35-day time-point (Fig. 1B, 1C; (*4*)), indicating that some mutant cells are never eliminated from the pancreas *in vivo*. Consistent with our previously published report (*4*), we found rare Alcian blue positive lesions (SFig. 1A, left panels) in 6/9 KrasG12D (KC) mice at 168 days p.i., suggesting some non-eliminated KrasG12D cells progress to pancreatic intraepithelial neoplasia (PanINs) (*25*). We did not detect PanIN lesions in p53R172H (PC) tissues. As expected, PanIN lesions were detected more frequently in 5/5 KPC tissues analysed (SFig. 1A, right panel) and 4/5 mice developed PDAC tumours by 168 days p.i.

We hypothesised that non-eliminated mutant cells override competitive cell elimination signals via common mechanisms. A deeper understanding of how KrasG12D cells avoid cell elimination may inform on the biology of early tumorigenesis. To gain an unbiased insight into how non-eliminated mutant cells survive in tissues we performed bulk RNA sequencing (Fig. 2A). Whole pancreata were harvested from experimental cohorts (KC/PC/KPC/Control) at 35 days p.i. and digested to cell suspensions. We used anti-lectin antibodies to enrich for acinar/ductal epithelial cells, as it is generally accepted that PDAC originates from genetically mutant acinar and/or ductal epithelial cells (*26*), and FACS sorted leptin positive RFP positive cells. To ensure we reached a minimum of 5,000 viable lectin positive RFP positive cells per sample, we pooled cell suspensions generated from three pancreas tissues per genotype. This was repeated to generate three samples per genotype for sequencing. We compared differential gene expression in non-eliminated mutant cells to wild type controls and found that KrasG12D cells (KC) predominantly upregulated genes, while p53R172H (PC) and double mutant (KrasG12D p53R172H, KPC) cells downregulated genes (Suppl. Table S1, S2). We found few differentially expressed genes in common between non-eliminated mutant cells when compared to wild type control (see Suppl. Table S1-S3).

**Figure 2:**
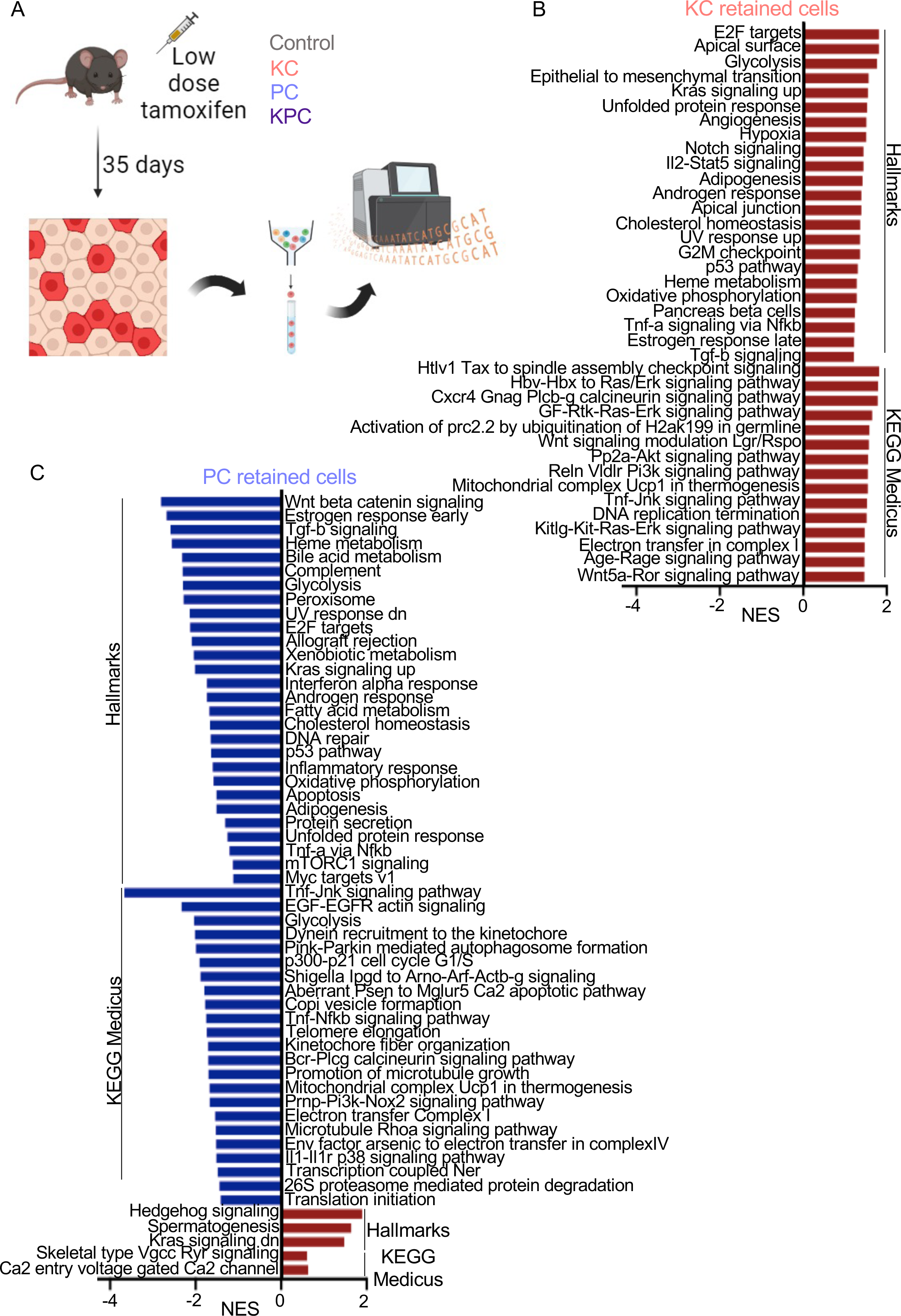
Bulk RNA sequencing results indicate cell fate changes in p53R172H and KrasG12D mutant retained cells. **A** Experimental design for tissue collection for RNA sequencing. Control, KC, PC and KPC mice were treated with low dose tamoxifen induction. RFP+ cells were isolated from pancreas tissues harvested at 35 days p.i. for bulk RNA sequencing. Three samples were analysed per genotype, each sample was obtained by pooling the pancreas from three mice. Illustration created with BioRender.com. **B-C** Normalised enrichment scores (NES) of gene set enrichment analysis (GSEA) on the Hallmark and KEGG MEDICUS gene sets for the RNA-seq analysis of **B** KC retained cells or **C** PC retained cells. Only gene sets with an FDR of less than 0.25 were included in the graph. The complete list that contains the results of GSEA analysis is provided in supplementary tables S4-S7.

To elucidate the biological pathways underpinning mutant cell retention in pancreas tissues, we applied gene set enrichment analysis (GSEA_4.2.2; entire Hallmarks and KEGG MEDICUS databases) and compared non-eliminated KrasG12D (KC) or p53R172H (PC) transcriptomes to wild type controls. We found that KrasG12D (KC) signatures positively correlated with MAPK (Fig. 2B, SFig. 2A) and p53 (Fig. 2B, SFig. 2B) signalling pathways, compared to controls, whereas p53R172H (PC) signatures negatively correlated with p53 (Fig. 2C, SFig. 2C) and with Kras (Fig. 2C) and MAPK (SFig. 2D) signalling. Non-eliminated KPC signatures also downregulated KRAS signalling (SFig. 2E), suggesting deregulation of Kras signalling and/or p53 pathway is a general requirement for mutant cells to remain in tissues. We found that KrasG12D (KC) signatures positively enriched for pathways associated with stress responses (Fig. 2B, e.g., Oxidative phosphorylation, Hypoxia, Unfolded protein response, UV response), deregulation of cell cycle (e.g., E2F targets and G2/M checkpoint) and immune response (e.g., Il2-Stat5 signalling, TNFa via NFkB signalling). In contrast, pathways associated with p53, cell cycle and inflammatory responses were negatively enriched in p53R172H (PC) signatures (Fig. 2C). Notably, KrasG12D (KC) signatures positively enriched for pro-survival and pro-tumorigenic pathways (Fig. 2B, e.g., Epithelial to Mesenchymal Transition (EMT), TNF-α signalling, Hypoxia, Angiogenesis, Notch signalling, Wnt signalling). Gene signatures from both p53R172H (PC; Fig. 2C) and KrasG12D p53R172H (KPC; SFig. 2E) negatively enriched for apoptosis, suggesting cell survival is also a requisite for mutant cell retention in tissues.

### Gene signatures of non-eliminated KrasG12D cells correlate with PDAC initiation and with increased stemness and cellular reprogramming

Mouse models of pancreatic cancer show that expression of oncogenic *Kras* triggers injury and stress responses, which often translate as cellular reprogramming and a change in cell fate (*27*). We found that KrasG12D (KC) gene signatures positively correlated with WP_Pancreatic_Adenocarcinoma_Pathway in GSEA analysis (SFig. 2F). Consistent with GSEA analyses, we found that gene signatures of non-eliminated PC (p53R172H) and KPC (KrasG12D p53R172H) cells negatively correlated with pro-tumorigenic pathways (Fig. 2C, SFig. 2E, 2G, 2H). Using published datasets (*27*), we screened transcriptomes of non-eliminated cells and found consistent upregulation of pancreas stem and progenitor genes (e.g., *Nkx6-1*, *Prom-1*, *Pdx-1, Hnf1a*) in non-eliminated KrasG12D (KC) populations (Fig. 3A). We also found canonical markers of spasmolytic polypeptide-expressing metaplasia (SPEM (*28*); e.g., *Aqp5, Tff2, Dmbt1, Gkn3, Wfdc2*) were upregulated in non-eliminated KrasG12D (KC) cells (SFig. 2I), as well as acinar genes (e.g., *Ptf1a, Rbpjl*), ductal genes (e.g., *Sox9, Onecut1*) and mucin genes (e.g., *Muc5a*, *Spdef, Creb3l4, Creb3l1*) (SFig. 2J). Finally, we found stemness gene signatures enriched in KrasG12D cells (Ramalho_stemness_up (M9473) and Malta_curated_stemness_markers (M30411) GSEA gene sets; KC) compared with control (SFig. 2K, 2L). We did not see enrichment of these gene signatures in p53R172H-expressing populations (SFig. 2I, 2J, 2M, 2N). We consistently showed that ductal epithelial compartments efficiently eliminate mutant cells and very few RFP+ cells remain in ducts at 35 days p.i. cells (Fig. 1D; (*4*)). Thus, we assume that most RFP+ cells isolated for RNA sequencing are acinar in origin. Gene signature data imply that non-eliminated KrasG12D populations represent differentiated acinar cells and early embryonic pancreatic progenitors, cells undergoing acinar-ductal metaplasia, gastric pyloric and intestinal metaplasia. The data also demonstrate that canonical KrasG12D-induced disease-associated metaplasia is evident in tissues at very early time points (35 days post activation of the transgene).

**Figure 3:**
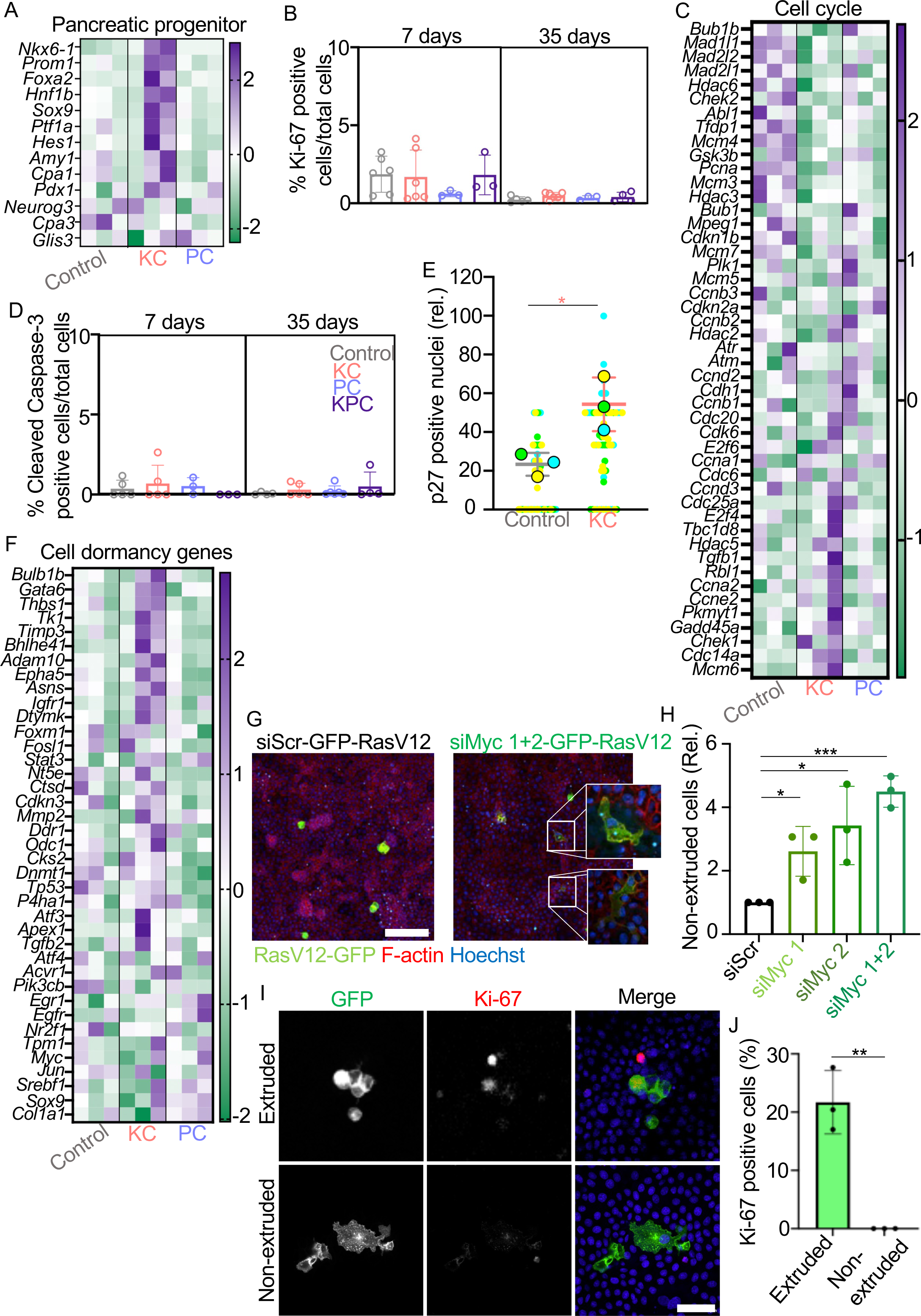
Cell dormancy/diapause gene signatures and cell cycle arrest are features of non-eliminated Ras mutant cells. Heatmaps showing row z-scores representing expression of **A** pancreatic progenitor associated, **C** cell cycle associated and **F** cell dormancy (rows) in non-eliminated KrasG12D (KC), non-eliminated p53R172H (PC) or Control cells (columns). Only genes with z-scores of a foldchange of 0.5 for KC average values are shown in Cell cycle panel. The full list of genes can be found in supplementary table S10. **B** Percentage of Ki-67 positive cells/total cells in Control, KrasG12D (KC), p53R172H (PC) or double mutant (KPC) pancreas tissues harvested at 7 and 35 days p.i. The data represent mean and +/- s.d. per mouse; control n=6 (7-days p.i.), n=5 (35-days p.i.); KC n=6 (7-and 35-days p.i.); PC n=3 (7-and 35-days p.i.); KPC n=3 (7-days p.i.), n=4 (35-days p.i.) mice. **D** Percentage of cleaved caspase-3 positive cells/total cells in Control, KrasG12D (KC), p53R172H (PC) or double mutant (KPC) pancreas tissues harvested at 7 and 35 days p.i. mean +/- s.d. per mouse; control n=5 (7-days p.i.), n=4 (35-days p.i.); KC n=5 (7- and 35-days p.i.); PC n=3 (7-days p.i.), n=6 (35-days p.i.); KPC n=3 (7-days p.i.), n=4 (35-days p.i.) mice. **E** Percentage of p27 positive nuclei per total number of nuclei in each RFP+ cluster in Control or KrasG12D (KC) pancreas tissues harvested at 35 days p.i. SuperPlot shows all the quantified RFP+ clusters (smaller circles) and the mean for each mouse (n=3 samples; larger circles - blue, green, and yellow). The graph shows mean and +/- s.d. for the three mice. Student’s t test was used to analyse the data, \**p*=0.0237. **G** Confocal images of GFP-RasV12 cells-expressing scrambled siRNA (siScr) or two siRNA oligos targeting endogenous *Myc* (siMyc1+2) and then mixed with non-labelled MDCK cells (1:50). Cells were stained with phalloidin to visualise F-actin (red) and Hoechst (blue). Merged images are shown. Scale bar, 100µm. **H** The proportion of non-extruded GFP-RasV12 cells transiently expressing different siRNA *Myc* oligos (siMyc1, siMyc2, siMyc1+2) relative to non-extruded GFP-RasV12 cells expressing siRNA scrambled (siScr). Data represent mean +/- s.d. from three independent experiments. At least 300 cells per point (100 cells per experiment). Student’s t test was used to analyse the data, ***p<0.0005, *p<0.05. **I** Confocal images of GFP-RasV12-expressing siRNA oligos targeting endogenous Myc (siMyc1+2), mixed with non-labelled MDCK cells (1:50). Cells were stained with anti-Ki-67 antibodies (red) and Hoechst (blue). Scale bar, 50 µm. **J** Percentage of Ki-67 positive cells extruded GFP-RasV12 cells as a fraction of total extruded cells. Non-extruded cells were not Ki-67 positive. Data represent mean +/- s.d. for 60 cells from three independent experiments. Student’s t test was used to analyse the data, **p<0.005.

### Non-eliminated KrasG12D mutant cells express features of cell dormancy

Several of our data indicated that non-eliminated cells are not actively dividing in tissues over time, e.g., we found no change in the amount of RFP fluorescence in tissues over time (Fig. 1C), suggesting RFP+ cells are not expanding. Consistently (*4*), immunostaining for Ki-67 showed an absence of proliferation in all tissues (Fig. 3B). GSEA analyses identified enrichment of proliferation-related pathways (e.g., E2F targets, GF-Rtk-Ras-Erk signalling) in non-eliminated KrasG12D cells compared to controls (KC; Fig. 2B); however, analysis of specific gene signature data showed a general downregulation of cell cycle genes (KC; Fig. 3C). GSEA analysis also revealed enrichment of pathways indicative of dysregulation of the cell cycle towards genomic instability (*29, 30*) (Fig. 2B, e.g., G2M checkpoint, Htlv Tax to spindle assembly checkpoint, Hbv-Hbx to Ras/Erk signalling, DNA-replication termination). Cell cycle gene expression profiles of non-eliminated p53R17H cells compared to wild type controls (PC; Fig. 3C) was like that of KrasG12D signatures and GSEA analyses indicated a general downregulation of proliferation-related pathways (e.g., E2F targets, Myc targets, Kras signalling up) in mutant p53-expressing cells (PC; Fig. 2C; KPC; SFig. 2E). Using β-galactosidase as a readout, we found no evidence of cell senescence in 35-day KrasG12D (KC) pancreas tissues (SFig. 3A). We also counted rare caspase-3 positive events in all pancreas tissues analysed (Fig. 3D), suggesting a general absence of apoptosis. We hypothesised that non-eliminated cells deregulate and/or arrest in the cell cycle. Using p27 (also known as Kip1) immunostaining as a readout of cell cycle arrest (*31*), we scored a significant increase in p27 positive RFP positive nuclei in KrasG12D (KC) tissues compared to controls (Fig. 3E; *p*=0.0237), suggesting non-eliminated KrasG12D cells are arrested at G_1_ stage of the cell cycle. Interestingly, non-eliminated KrasG12D (KC) gene signatures were also enriched for pathways associated with cancer cell dormancy, (e.g., Oxidative phosphorylation, Unfolded protein response, Hypoxia, Fig. 2B) (*32*). In general, cell dormancy describes a reversible cell cycle arrested state, often triggered by cell stress (*33–35*). Using published gene signatures of cell dormancy as reference (*36*), we found cell dormancy (Fig. 3F) and diapause (SFig. 3B) gene signatures are enriched in non-eliminated KrasG12D (KC) cells compared to control cells. Diapause is a temporary halt in embryogenesis when conditions are detrimental to development (*35*). We also found NRF2 pathway (SFig. 3C) and NRF2 target genes (SFig. 3D) were positively enriched in non-eliminated KrasG12D (KC) cells. NRF2 is a transcription factor, which is activated during cell dormancy (*37*). Cell dormancy/diapause gene signatures or related pathways were not significantly enriched in non-eliminated p53R172H (PC) cells (Fig 3F, SFig. 3B). These data infer that non-eliminated mutant cells deregulate the cell cycle towards an arrested state; however, two copies of functional p53 are required for cells to adopt a dormant or diapaused state (*38–40*).

### Cell cycle arrest prevents mutant cell extrusion *in vitro*

We hypothesised that mutant cells override cell elimination *in vivo* by arresting in the cell cycle. Indeed, extrusion of v-Src-expressing cells from normal epithelial sheets is dependent on the cell cycle (*41*). To functionally test the requirement for cell cycle arrest in RasV12 cell extrusion, we used previously established cell competition assays and Madin-Darby Canine Kidney (MDCK) cells expressing GFP-tagged constitutively active oncogenic RasV12 (GFP-RasV12) (*5, 9*). We have previously shown that when surrounded by normal MDCK cells, GFP-RasV12-expressing MDCK cells are outcompeted and eliminated from an epithelial sheet via cell extrusion (*5, 9*). Once extruded, GFP-RasV12 cells rapidly proliferate to form multicellular aggregates (*5*). In contrast, MDCK cells expressing mutant p53 (p53R175H, p53R273H) are outcompeted by normal cells via necroptosis (*42*). We induced cell cycle arrest in GFP-RasV12 cells via *Myc* knockdown and transient transfection of two independent c-Myc siRNAs (transfected alone or in combination; siMyc-GFP-RasV12). We chose *c-Myc* knockdown as an approach because c-Myc is a key regulator of the cell cycle (*43*) and depletion of Myc triggers cell dormancy (*44*). Expression of c-Myc siRNA yielded a decrease in MYC protein levels, 48h post-transfection (SFig. 3E, 3F), an increase in p21 protein expression and decreased phospho-p38 protein levels (SFig. 3E, 3F) and a marked reduction in GFP-RasV12 cell confluency (SFig. 3G) compared to GFP-RasV12 cells expressing scrambled siRNA controls (siScr-GFP-RasV12). Thus, transient depletion of Myc in GFP-RasV12 cells promotes cell cycle arrest. To assess the impact of cell cycle arrest on normal-mutant cell-cell interactions, we used our established cell extrusion assay (*5, 9*), whereby GFP-RasV12 cells were first transfected with siScr or siMyc oligos and then mixed with normal MDCK cells at 1:50 ratios. At 48h post mixing, cells were fixed, stained with phalloidin and Hoechst to visualise F-actin and nuclei respectively. We scored apical extrusion events and found that the proportion of non-extruded GFP-RasV12 cells significantly increased when GFP-RasV12 cells were depleted for MYC (siMyc-GFP-RasV12; Fig. 3G, 3H; *p*=0.0003 siMyc1+2-GFP-RasV12 compared to siScr-GFP-RasV12 controls). Immunofluorescence staining using anti-Ki-67 antibodies confirmed that non-extruded siMyc-GFP-RasV12 cells are not proliferating, whereas the minority of Ki-67 positive cells were extruded (Fig. 3I, 3J). We also used cell confrontation assays (*9*) and live-cell imaging of normal-mutant interactions across an entire epithelial cell sheet (SFig. 3H). We previously reported that upon collision with normal MDCK cells, GFP-RasV12 cells are triggered to retract and segregate away from normal cells, separating via the formation of smooth boundaries (*9*). We found that GFP-RasV12 cells depleted for MYC (siMyc1+2) retracted less efficiently than GFP-RasV12 siScr controls (SFig. 3I, SFig.3J; p=0.0086). Thus, RasV12 cells depleted for Myc and exhibiting cell cycle arrest are not extruded or triggered to segregate by normal cells, suggesting cell cycle arrest protects RasV12 mutant cells from cell elimination.

### Non-eliminated KrasG12D cells activate β-catenin independent Wnt signalling *in vivo*

The Wnt pathway positively regulates cancer cell dormancy (*45, 46*), diapause (*32*) and stemness (*47, 48*) in different tissue contexts. Our GSEA analyses identified an enrichment for Wnt pathway activation in non-eliminated KrasG12D (KC) transcriptomes compared to controls (Fig. 2B, Wnt signalling modulation Lgr/Rspo, Wnt5a-Ror signalling pathway) and increased gene expression of both Wnt target genes and components of the Wnt pathway such as ligands and receptors (Fig. 4A). Genes commonly associated with the Wnt/β-catenin pathway were generally downregulated in KrasG12D (KC) cells, including *Lgr5*, *Myc*, *Sox2*, *Nanog*, *Esrrb* and *Wnt3a* (Fig. 4B). Similarly, GSEA analyses indicated low levels of Wnt/β-catenin signalling in non-eliminated p53R172H (PC) cells (Fig. 2C), which is supported by a general downregulation of Wnt pathway genes (SFig. 4A). Interestingly, both KrasG12D (KC; SFig. 4B) and p53R172H (PC; SFig. 4C) gene signatures positively correlated with Wnt5a-Ror signalling pathway. Wnt ligands and receptors associated with Wnt5a-Ror signalling were increased in KrasG12D (KC) cells, including *Wnt5a*, *Ror1*, *Ror2*, *Dvl2* and *Jun* (Fig. 4C, 4D and Supplementary table S11).

**Figure 4:**
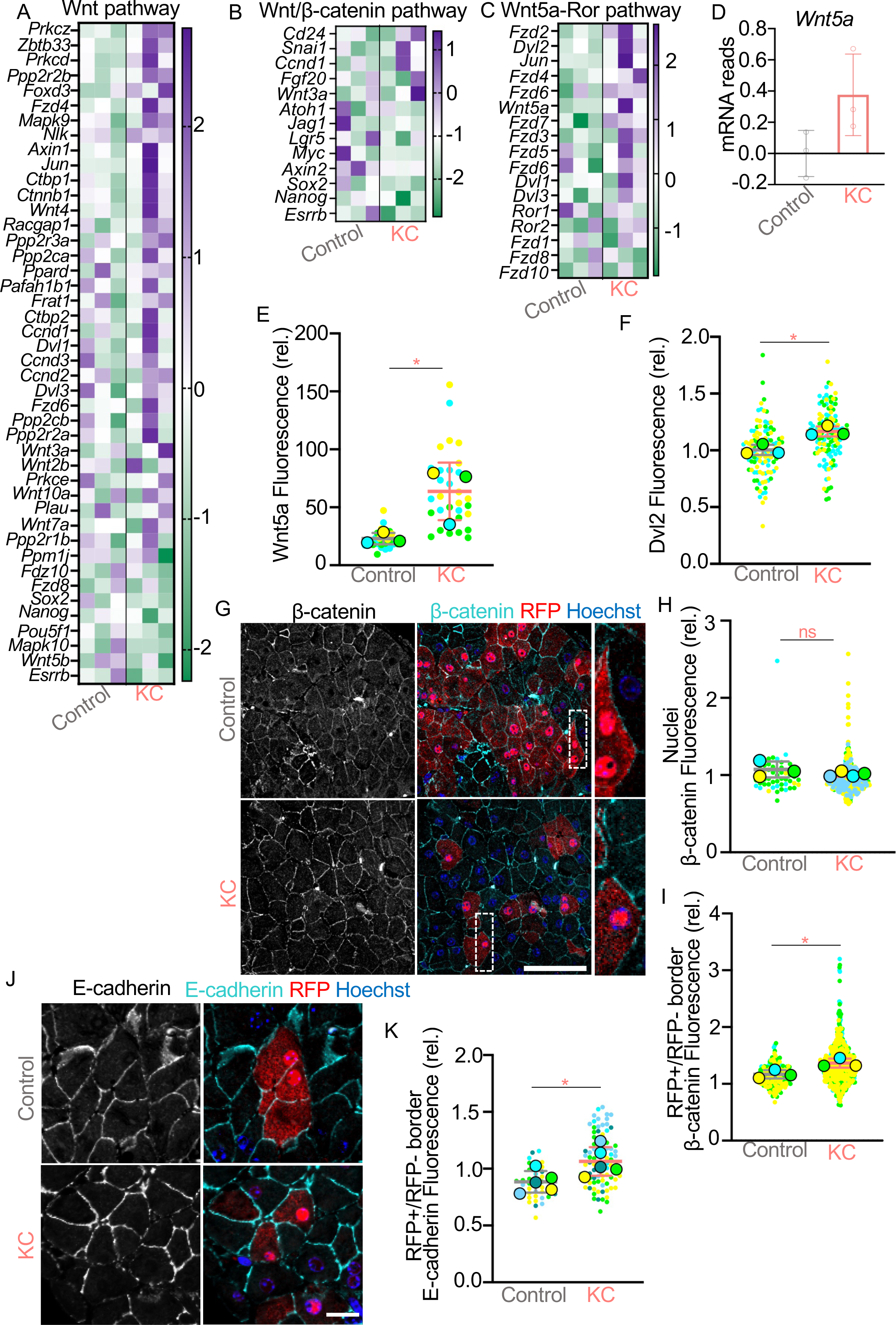
Wnt pathway is activated at normal-mutant cell-cell boundaries *in vivo*. Heatmaps of **A** Wnt signalling pathway, **B** Wnt/β-catenin pathway or **C** Wnt5a-Ror pathway related genes (rows) in Control and KrasG12D (KC) cell transcriptomes (columns). The heatmaps show row z-scores for each gene obtained from the RNA sequencing experiment. Data from three samples (three mice pooled per sample) per genotype are shown. Only genes with z-scores of a foldchange of 2 are shown in the Wnt pathway panel. The full list of genes can be found in supplementary table S11. **D** *Wnt5a* mRNA reads relative to control reads in the three samples in Control and three samples in KrasG12D (KC) obtained from the RNA sequencing experiment. Graph represents mean +/- s.d. reads. **E** Mean fluorescence intensity of WNT5A in Control or KrasG12D (KC) pancreas tissues harvested at 35 days p.i. SuperPlot shows all the quantified tissue areas (smaller circles) and the mean for each mouse (n=3 samples; larger circles - blue, green, and yellow). The graph shows mean and +/- s.d. for the three mice. Student’s t test was used to analyse the data, \**p*=0.0491. **F** Mean fluorescence intensity of DVL2 in Control or KrasG12D (KC) pancreas tissues harvested at 35 days p.i. Mean fluorescence intensity is relative to the background. SuperPlot shows all the quantified RFP+ clusters (smaller circles) and the mean for each mouse (n=3 samples; larger circles - blue, green, and yellow). The graph shows mean and +/- s.d. for the three mice. Student’s t test was used to analyse the data, \**p*=0.0110. **G** Confocal images of pancreas tissues harvested at 35-days p.i. from Control, KrasG12D (KC) mice. Tissues were stained with anti-RFP (red) and anti-β-catenin (cyan) antibodies and Hoechst (blue). Dashed boxes represent digital zoom images of representative cell-cell contacts. Scale bar, 50µm. **H** Mean fluorescence intensity of β-catenin in the nuclei of RFP positive cells in Control and KrasG12D (KC) tissues harvested at 35 days p.i. Mean fluorescence intensity is relative to the background. SuperPlot shows all the quantified nuclei (smaller circles) and the mean for each mouse (n=3 samples; larger circles - blue, green, and yellow). The graph shows mean and +/- s.d. for the three mice. Student’s t test was used to analyse the data. ns p=0.3035. **I** Mean fluorescence intensity of β-catenin at the boundary between RFP positive and negative cells in Control and KrasG12D (KC) tissues harvested at 35 days p.i. Mean fluorescence intensity is relative to the background. Fluorescence was measured in ROIs, separated by a constant distance, along the cell-cell boundary. SuperPlot shows all the quantified ROIs (smaller circles) and the mean for each mouse (n=3 samples; larger circles - blue, green, and yellow). The graph shows mean and +/- s.d. for the three mice. Student’s t test was used to analyse the data. *p=0.0347. **J** Confocal images of pancreas tissues harvested at 35-days p.i. from Control and KrasG12D (KC) mice and stained with anti-RFP (red) and anti-E-cadherin (cyan) antibodies and Hoechst (blue). Scale bar, 20µm. **K** Mean fluorescence intensity of E-cadherin at the boundary between RFP positive and RFP negative cells in Control and KrasG12D (KC) tissues harvested at 35 days p.i. E-cadherin fluorescence was quantified and reported as in I. SuperPlot shows all the quantified ROIs (smaller circles) and the mean for each mouse (n=5 samples; larger circles – lighter blue, blue, darker blue, green, and yellow). The graph shows mean and +/- s.d. for the five mice. Student’s t test was used to analyse the data. *p=0.0348.

To investigate active Wnt signalling *in vivo*, we first performed immunofluorescence for CD44 which is a target of β-catenin dependant and independent signalling (*49–51*). Consistent with previous reports (46), we observed CD44 labelling at cell membranes in PanINs (SFig. 4D). We found that CD44 was significantly increased at RFP+: RFP-cell-cell boundaries in KrasG12D (KC) tissues compared to RFP+: RFP-boundaries in wild type controls (SFig. 4E, 4F; KC versus control, *p*=0.0009). In contrast, CD44 was significantly reduced within a cluster of KrasG12D cells (KC; SFig. 4E, 4G; *p*=0.0455). We asked whether increased CD44 at mutant-normal cell-cell boundaries was dependent on competitive cell-cell interactions. We quantified CD44 fluorescence at cell-cell contacts between RFP+/RFP-cells in KC tissues fixed following high dose tamoxifen induction (KC Megadose). We previously reported that KrasG12D cells are not eliminated from megadose tissues, where mutant cells occupy ∼80% of the tissue (*4*). We found that CD44 levels were significantly lower at mutant-normal cell boundaries in megadose tissues compared to KrasG12D low dose tissues (KC versus KC Megadose; SFig 4E, 4F; *p*=0.0047), while CD44 levels were significantly higher between KrasG12D cells in megadose tissues (SFig. 4E, 4G; *p*=0.0341). Similarly, we found that CD44 was significantly increased at RFP+: RFP-cell-cell boundaries in p53R172H tissues (PC) compared to RFP+: RFP-boundaries in wild type controls (SFig. 4H; PC versus control, *p*=0.047). CD44 was significantly reduced within a cluster of p53R172H cells (SFig. 4I, PC versus control, *p=*0.0397). We next asked whether readouts of Wnt5a signalling were evident in non-eliminated KrasG12D cells in fixed pancreas tissues. Using quantitative immunofluorescence protocols and commercial antibodies, we observed significant increase in WNT5A (Fig. 4E; *p=*0.0491) and DVL2 (Fig. 4F; *p=*0.0110) protein in KrasG12D (KC) cells relative to controls, suggesting Wnt5a signalling is active in non-eliminated KrasG12D cells *in vivo*. In addition, we performed immunofluorescence for β-catenin in tissues fixed at 35 days p.i., using nuclear localisation as a surrogate marker of activity of the Wnt/ β-catenin signalling pathway. Nuclear β-catenin was undetectable in nuclei of both control and KrasG12D (KC) cells (Fig. 4G, 4H; *p*=0.3035), indicating that Wnt/β-catenin signalling is not active in non-eliminated KC cells, consistent with our transcriptional analysis. Instead, β-catenin was significantly increased at RFP+: RFP-cell-cell contacts in KrasG12D (KC) tissues compared to controls (Fig. 4G, 4I; *p*=0.0347). Moreover, levels of E-cadherin were also significantly elevated at RFP+: RFP-boundaries in KrasG12D (KC) tissues compared to controls (Fig. 4J, 4K; *p*=0.0348). Interestingly, we also detected increased mRNA expression of atypical *cadherin-6* in non-eliminated KrasG12D cell signatures (KC; SFig. 5A) and increased levels of CADHERIN-6 (K-cadherin) in RFP+ clusters in KrasG12D (KC) tissues only (SFig. 5B, 5C; *p*=0.0474). Together, our data suggest that β-catenin-independent Wnt signalling is active, specifically in non-eliminated KrasG12D cells and at KrasG12D-normal cell-cell contacts *in vivo*. Our data also infer that active Wnt signalling increases cohesiveness and/or stability of E-cadherin-based cell-cell adhesions *in vivo*, specifically at cell-cell boundaries between non-eliminated mutant cells and normal neighbours.

### Wnt5a inhibits RasV12 cell extrusion *in vitro* by stabilising E-cadherin-based cell-cell adhesions

To establish whether Wnt5a directly inhibits RAS mutant cell elimination, we returned to MDCK cell extrusion assays (*5, 9*). We first assessed the effect of Wnt5a treatment on normal-RasV12 cell-cell interactions. We found that the proportion of non-extruded GFP-RasV12 cells significantly increased in the presence of Wnt5a (Fig. 5A, right panel, yellow arrowheads; 5B, yellow bar; *p*=0.0065) compared to PBS treated controls (Fig. 5A, left panel, white arrowheads; 5B, black bar). Remarkably, addition of Wnt3a had no effect on RasV12 cell extrusion rates compared to PBS controls (Fig. 5A, middle panel, white arrowheads; 5B, pink bar; *p*=0.4488). In general, Wnt3a and Wnt5a activate the β-catenin-dependent and β-catenin-independent pathway respectively (*52, 53*), suggesting β-catenin-independent Wnt signalling inhibits RasV12 cell extrusion.

**Figure 5:**
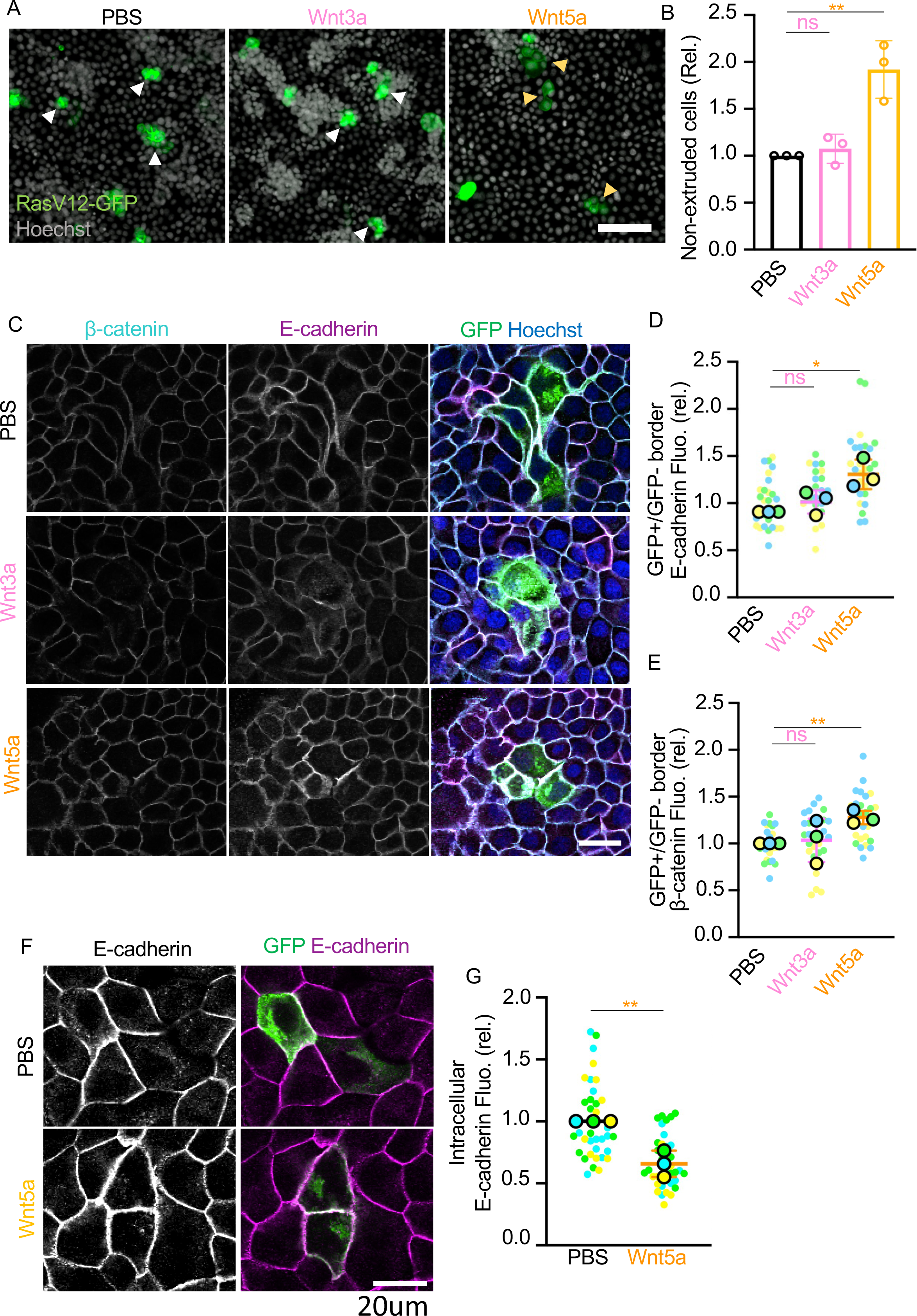
Wnt5a induces stable E-cadherin-based cell-cell adhesion in RasV12 cells. **A** Confocal images of extruded and non-extruded cells treated with PBS, Wnt3a or Wnt5a. Fixed cells were stained with Hoechst (grey). Arrowheads indicate extruding (white) or non-extruding (yellow) GFP-RasV12 cells. Scale bar, 100 µm. **B** The proportion of non-extruded GFP-RasV12 cells treated with Wnt3a (pink) or Wnt5a (yellow) recombinant proteins relative to non-extruded GFP-RasV12 cells treated with PBS (black). The data represent mean +/- s.d. for at least 300 cells from three independent repeats. Student’s t test was used to analyse the data. **p=0.0065; ns p=0.4488. **C** Confocal images of GFP-RasV12 cells mixed with MDCK cells treated with PBS, Wnt3a or Wnt5a for 30h. Cells were stained with anti-β-catenin (cyan) and anti-E-cadherin (magenta) antibodies and Hoechst (blue). Scale bar, 20µm. **D, E** Mean fluorescence intensity of E-cadherin (**D**) or β-catenin (**E**) at the boundary between GFP-RasV12 and MDCK cells treated with PBS (black), Wnt3a (pink) or Wnt5a (yellow) for 30h. Fluorescence was measured in ROIs, separated by a constant distance, along the cell-cell boundary. SuperPlot shows all the quantified ROIs (smaller circles) and the mean for each experiment (n=3 samples; larger circles - blue, green, and yellow). The graph shows mean and +/- s.d. for the three experiments. Student’s t test was used to analyse the data. **p=0.0026; *p=0.0116; ns p>0.2. **F** Confocal images of GFP-RasV12 cells mixed with MDCK cells treated with PBS or Wnt5a for 16h. Cells were stained with anti-E-cadherin (magenta) antibody and Hoechst (blue). Scale bar, 20µm. **G** Mean fluorescence intensity of intracellular E-cadherin in GFP-RasV12 (mixed with MDCK cells) treated with PBS (black) or Wnt5a (yellow) for 16h. SuperPlot shows all the quantified GFP-RasV12 cells (smaller circles) and the mean for each experiment (n=3 samples; larger circles - blue, green, and yellow). The graph shows mean and +/- s.d. for the three experiments. Student’s t test was used to analyse the data. *p=0.0051.

Cell extrusion requires dynamic remodelling of E-cadherin-based cell-cell adhesions at normal-mutant cell-cell boundaries (*5, 54, 55*). Considering both E-cadherin and β-catenin protein were significantly increased at normal-mutant cell-cell boundaries *in vivo* (Fig. 4G, 4I-4K), we asked whether treatment with exogenous Wnt ligands modulates adherens junctions at normal-mutant cell-cell boundaries in MDCK assays. We analysed E-cadherin and β-catenin by immunofluorescence in Wnt-treated co-cultures and focused on cell-cell interactions between GFP-RasV12 that had not yet been extruded and direct normal neighbours. In the presence of Wnt3a, β-catenin and E-cadherin staining appeared diffuse/weak and sometimes absent at normal-mutant cell-cell contacts (Fig. 5C, middle panels), similar to PBS-treated controls (Fig. 5C, top panels). In contrast, E-cadherin and β-catenin staining was more defined at normal-mutant cell-cell contacts in Wnt5a treated cells (Fig. 5C, lower panels). Quantitative analysis showed that in Wnt5a treated cells, E-cadherin (Fig. 5D; *p*=0.0116) and β-catenin (Fig. 5E; *p*=0.0026) significantly increased at normal-RasV12 cell-cell contacts. In contrast, Wnt3a treatment had no significant effect on E-cadherin (Fig. 5D; *p*=0.2137) or on β-catenin (Fig. 5E; *p*=0.8111) at RasV12-normal cell-cell contacts when compared to PBS-treated controls. Treatment with Wnt5a significantly increased the levels of E-cadherin (SFig. 5D; *p*=0.0003) but had no effect on β-catenin levels (SFig. 5E; *p*=0.6049) detected at cell-cell contacts between RasV12 cells, suggesting Wnt5a signalling specifically modulates E-cadherin at cell-cell contacts. Consistently Wnt3a had no significant effect on E-cadherin or β-catenin at cell-cell contacts between RasV12 cells (SFig. 5D; *p*=0.0584; SFig. 5E; *p*=0.2828). Moreover, in cell confrontations assays (*9*), Wnt5a treatment significantly reduced RasV12 cell speed (SFig. 5F; *p*=0.0268) compared to PBS treated controls, whereas normal MDCK cell speed was unaffected (SFig. 5F; *p*=0.8202), implying Wnt5a signalling specifically affects RasV12 cell cohesion.

E-cadherin endocytosis is required for apical extrusion of RasV12 cells from normal MDCK cell sheets (*55*); therefore, we asked whether Wnt5a abrogates this process to prevent cell extrusion. We imaged RasV12-normal cell-cell contacts at 16h (prior to cell extrusion events (*55*)) using super resolution microscopy. At this time point, E-cadherin staining was weakly visible/diffuse at RasV12-normal cell-cell contacts in PBS-treated controls and was often detected in distinct intracellular puncta in both RasV12 and normal cells (Fig. 5F, top panels). In contrast, Wnt5a treated cells showed strong, defined E-cadherin at cell-cell contacts and E-cadherin positive puncta were less visible in RasV12 cells and neighbouring cells (Fig. 5F, lower panels). Quantification of fluorescence intensity of intracellular E-cadherin showed a significant reduction in RasV12-cells treated with Wnt5a compared to PBS-treated controls (Fig. 5G; *p*=0.0051), suggesting Wnt5a treatment blocks E-cadherin recycling/endocytosis at RasV12-normal boundaries. Finally, we immunostained for Caveolin-1 (CAV1) a key regulator of endocytosis (*56*) and cell extrusion (*57, 58*). CAV1 was detected in normal MDCK and RasV12 cells at cell-cell junctions (SFig. 5G). At the timepoint when endocytosis is evident in cells (Fig. 5F), CAV1 fluorescence was significantly reduced at RasV12-normal cell-cell contacts in Wnt5a treated cells compared to PBS controls (SFig. 5H; *p*=0.0117). Taken together our data show that Wnt5a stabilises E-cadherin-based cell-cell adhesion at normal-mutant boundaries potentially by preventing E-cadherin internalisation. Increased E-cadherin-based cell-cell adhesion correlates with a significant reduction in RasV12 cell extrusion and increase in RasV12 cell cohesion.

### Wnt signalling is required to promote retention of mutant cells *in vitro* and *in vivo*

Next, we set out to test whether Wnt signalling is required to prevent mutant cell elimination. We first repeated cell extrusion experiments using siScr-GFP-RasV12 or siMyc 1+2-GFP-RasV12 MDCK cells mixed with normal MDCK cells at 1:50 ratios, allowed cells to form cell-cell interactions and then treated with the porcupine inhibitor WNT-974 (or DMSO). Porcupine is an acyltransferase enzyme required for the lipidation and trafficking of Wnt proteins, which is a necessary step for ligand secretion (*59*). By inhibiting porcupine, Wnt ligands cannot be secreted and therefore cannot signal to cells. In MDCK cells, overexpression of RasV12 induces expression and secretion of Wnt5a in a porcupine-dependent manner (*60*), suggesting porcupine inhibition in our experiments blocks secretion of endogenous Wnt, including Wnt5a. The proportion of non-extruded siMyc 1+2-FP-RasV12 cells significantly reduced in the presence of WNT-974 (Fig. 6A, 6B; *p*<0.0005), compared to siScr-GFP-RasV12 treated cultures (Fig. 6A; *p*=0.6912). In cell confrontation assays, we found that WNT-974-treated siMyc 1+2-GFP-RasV12 cells efficiently retracted (SFig. 6A, 6B) and segregated from normal cells, compared to DMSO-treated siMyc 1+2-GFP-RasV12 cells. Quantification of the linearity of the boundary between colliding sheets of cells (*9*) confirmed this observation (SFig. 6C; WNT-974-treated cells: *p*=0.0020).

**Figure 6:**
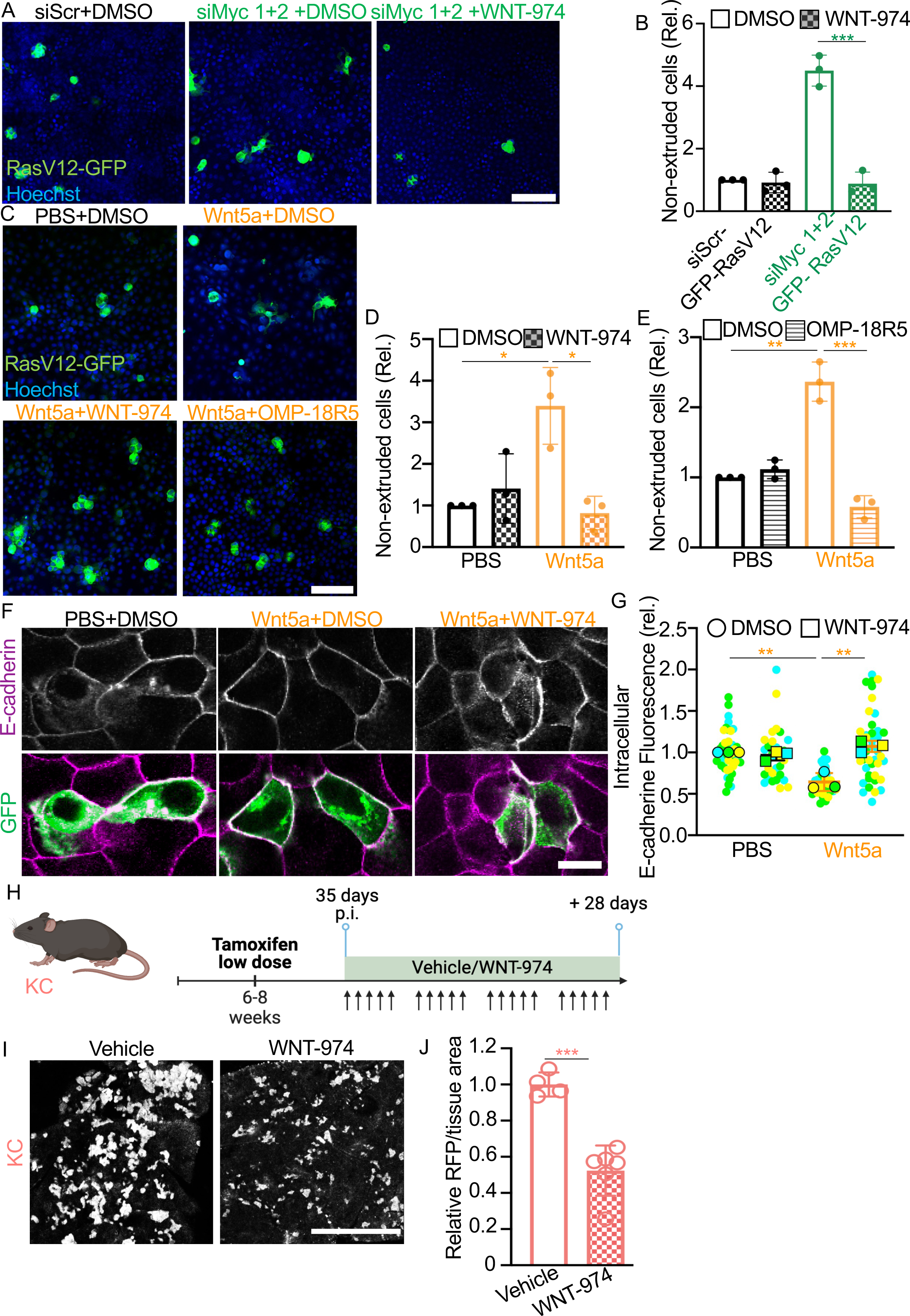
Wnt signalling prevents apical extrusion of RasV12 cells from normal epithelial monolayer *in vitro*. Inhibition of Wnt promotes elimination of non-eliminated cells *in vivo*. **A** Confocal images of extruded and non-extruded siScr-GFP-RasV12 and siMyc 1+2-GFP-RasV12 cells for 48h with DMSO or WNT-974. Fixed cells were stained with Hoechst (blue). Scale bar, 100 µm. **B** The proportion of non-extruded GFP-RasV12 cells transiently expressing siRNA *Myc* oligos (siMyc1+2) relative to non-extruded GFP-RasV12 cells expressing siRNA scrambled (siScr). Cells were treated with DMSO (white bars) or WNT-974 porcupine inhibitor (dashed bars). The data represent mean +/- s.d. for at least 300 cells from three independent repeats. Student’s t test was used to analyse the data. ***p<0.0005. **C** Confocal images of extruded and non-extruded GFP-RasV12 cells pre-treated with PBS/Wnt5a and treated for 48h with DMSO/WNT-974/OMP-18R5. OMP-18R5 is a Frizzled receptor antagonist. Fixed cells were stained with Hoechst (blue). Scale bar, 100 µm. **D, E.** GFP-RasV12 non-extruded cells pre-treated with PBS or Wnt5a recombinant protein 24h before mixing with MDCK cells (1:50). Six hours after mixing and plating, cells were treated with **D** DMSO or WNT-974 porcupine inhibitor or **E** OMP-18R5. The data represent mean +/- s.d. for at least 300 cells from three independent repeats. Student’s t test was used to analyse the data, ***p<0.0005, **p<0.005, *p<0.05. **F** Confocal images of GFP-RasV12 cells mixed with MDCK cells treated with PBS+DMSO, Wnt5a+DMSO or Wnt5a+WNT-974 for 16h. Cells were stained with anti-E-cadherin (magenta) antibody. Scale bar, 20µm. **G** Mean fluorescence intensity of intracellular E-cadherin in GFP-RasV12 (mixed with MDCK cells) treated with PBS (black) or Wnt5a (yellow) in combination with DMSO (circle) or WNT-974 (square) for 16h. SuperPlot shows all the quantified GFP-RasV12 cells (smaller circles) and the mean for each experiment (n=3 samples; larger circles or squares - blue, green, and yellow). The graph shows mean and +/- s.d. for the three experiments. Student’s t test was used to analyse the data. **p<0.005. **H** Experimental design for Wnt inhibition experiments *in vivo*. 6–8-week-old KC (*Kras^G12D/+^*) mice were induced with low-dose tamoxifen. Thirty-five days later, mice were treated with the porcupine inhibitor WNT-974 or vehicle for 28 days. Illustration created with BioRender.com. **I** Representative images of endogenous RFP fluorescence in KC pancreas tissue sections harvested 28-days post treatment with vehicle or WNT-974. Scale bar, 500µm. **J** RFP fluorescence per tissue area in KC mice treated with WNT-974 or vehicle and relative to vehicle treated tissues. Data represent mean per mouse (n=4-6 samples) and +/- s.d. Student’s t test was used to analyse the data, ***p=0.0003.

Consistently, the effects of Wnt5a on RasV12 cell extrusion rates were reversed in the presence of porcupine inhibitor (WNT-974; Fig. 6C, 6D, hatched bars; *p*=0.0114) and in the presence of the Frizzled (Fzd) receptor antagonist OMP-18R5 (*61*) (Fig. 6C, 6E, striped bars; *p*=0.0004) compared to DMSO-treated controls, resulting in a significant decrease in non-extruded GFP-RasV12 cells. Treatment with WNT-974 inhibitor also reversed Wnt5a-induced effects on E-cadherin localisation (Fig. 6F). As expected, Wnt5a induced a significant decrease in the level of intracellular E-cadherin fluorescence in RasV12 cells (Fig. 6G, circles; *p*=0.0048). In the presence of WNT-974 inhibitor, intracellular E-cadherin fluorescence significantly increased (Fig. 6G, squares; *p*=0.0044 compared to Wnt5a-treated cells) and returned to levels comparable to PBS treated controls, implying that Wnt signalling is required to prevent E-cadherin internalisation and endocytosis. In cell confrontation assays, Wnt5a treatment significantly decreased RasV12 cell speed (SFig. 6D, open bar; *p*=0.0156); RasV12 cell speed significantly increased in the presence of WNT-974 inhibitor (SFig. 6D, hatched bar; *p*=0.0058).

Finally, we set out to functionally test whether active Wnt signalling is necessary for mutant cell retention *in vivo*. Using 6-8-week-old adult KrasG12D (KC) mice, we first induced low level recombination in tissues, waited for 35 days p.i. to allow for cell competition to proceed and for most mutant cells to be eliminated, and then administered the porcupine inhibitor WNT-974 (or DMSO in corn oil vehicle) for a period of 4 weeks (Fig. 6H). To validate Wnt inhibition in tissues *in vivo*, we quantified CD44 fluorescence at mutant-normal cell boundaries and within RFP+ clusters in KrasG12D (KC) pancreas tissues treated with WNT-974 or vehicle. We found that levels of CD44 fluorescence significantly decreased in WNT-974 treated tissues compared to vehicle (SFig. 6E, *p*=0.0467), suggesting Wnt activity is reduced following treatment with porcupine inhibitor. Using global RFP fluorescence per tissue area as a readout of mutant cell fate, we measured a significant decrease in RFP fluorescence in the WNT-974-treated KrasG12D (KC) tissues compared to vehicle-treated animals (Fig. 6J, 6I; *p*=0.0003). We also found that porcupine inhibition induced a significant decrease in RFP fluorescence in p53R172H (PC) tissues compared to vehicle treated controls (SFig. 6F, 6G; *p*=<0.0001). Thus, inhibition of Wnt pathway induces the expulsion of KrasG12D or p53R172H cells from pancreas tissues *in vivo*; however, Wnt inhibition is not sufficient to remove all mutant cells, suggesting Wnt-independent mechanisms are also required. Nonetheless, our data support a model whereby activation of Wnt signalling allows mutant cells to avoid cell expulsion and to survive in the pancreas.

## Discussion

Tissue homeostasis is fundamental to healthy aging of an organism and cell competition is an important regulator of tissue health. Here, we extend our previously published results (*4*) to show that like KrasG12D-expressing cells, genetically mutant p53R172H-expressing cells compete with normal cells for survival in the adult pancreas and are often expelled from the tissue. We report that cell expulsion is inefficient, and a proportion of KrasG12D- or p53R172H-expressing cells are never eliminated from the tissue, suggesting these cells have a survival advantage. Transcriptomic analyses of non-eliminated mutant cells revealed Wnt signalling as a key regulator of mutant cell fate *in vivo*. Gene expression data indicate Wnt/β-catenin signalling is inactive and instead, non-canonical Wnt5a signalling is active in non-eliminated mutant cells. Indeed, readouts of non-canonical Wnt signalling are increased in non-eliminated KrasG12D cells and at KrasG12D-normal cell-cell boundaries *in vivo*. Wnt5a signalling (and not β-catenin-dependent Wnt3a signalling) inhibited apical cell extrusion of RasV12 cells from epithelial cell sheets *in vitro*, and this effect was reversed upon inhibition of porcupine/Fzd activity. In pancreas tissues, inhibition of Wnt signalling promoted expulsion of non-eliminated mutant cells, suggesting Wnt signalling is a general mechanism activated by mutant cells to override cell elimination mechanisms in the pancreas *in vivo*.

Apical cell extrusion is an active process that requires direct interaction with normal cells to expel genetically mutant cells from an epithelium (*62*). Here, we show that Wnt5a blocks apical extrusion of RasV12 cells, allowing mutant cells to remain in the epithelium. Mechanistically, we propose that Wnt5a stabilises E-cadherin-based cell-cell adhesions at RasV12-normal cell-cell contacts, and this prevents RasV12 cell extrusion. E-cadherin is a key modulator of apical cell extrusion: RasV12 cells are not extruded when E-cadherin is depleted in normal neighbours (*5*), suggesting E-cadherin-based cell-cell adhesion is required for RasV12 cell extrusion. On the other hand, our data support previous reports (*55*) that E-cadherin is internalised as discrete puncta at RasV12-normal cell-cell boundaries, suggesting E-cadherin must be dynamically remodelled for cell extrusion to occur. Indeed, inhibition of Rab5-mediated endocytosis prevents apical extrusion of RasV12 cells from MDCK cell sheets (*55*). Here, we show that in the presence of Wnt5a, E-cadherin puncta are not detected in mutant cells; instead, the levels of E-cadherin and β-catenin are increased at RasV12-normal cell-cell contacts. Moreover, inhibition of Wnt signalling (via porcupine inhibitor or Fzd antagonism) restores appearance of E-cadherin puncta at RasV12-normal cell-cell contacts and apical extrusion of RasV12 cells. In pancreas tissues, we previously showed that E-cadherin is decreased at KrasG12D-normal cell-cell adhesions, specifically at timepoints prior to KrasG12D cell elimination (*4*). Here, we extend these findings to show that E-cadherin and β-catenin are increased at cell-cell contacts between non-eliminated KrasG12D cells and normal neighbours. Indeed, we have shown that under conditions where RAS mutant cell elimination is prevented, E-cadherin increases at normal-mutant boundaries (*4*). Together our data support a model whereby stable cell-cell adhesion induced by Wnt5a signalling, prevents cell extrusion. Interestingly, we also found that caveolin-1 (CAV1), an important driver of RasV12 cell extrusion (*57, 58*) is also decreased at RasV12-normal cell-cell contacts following Wnt5a treatment. In development, Wnt5a signalling regulates cell cohesion and tissue fluidity via caveolin-dependent and clathrin-dependent endocytosis (*63*). Whether Wnt5a directly modulates endocytosis of E-cadherin at RasV12-normal cell-cell boundaries requires further investigation. Similar to our results, in human epithelial cells Wnt5a induces β-catenin membrane localisation and association with E-cadherin increasing intercellular adhesion (*64*).

We also show that RasV12 cells in a cell cycle arrested state were not extruded from MDCK monolayers. In pancreas, non-eliminated KrasG12D cells stained positive for p27, and upregulated cell dormancy signatures and pathways known to induce cell dormancy (e.g., Unfolded protein response, Wnt5a-ROR signalling, Hypoxia) (*65*). Thus, exiting the cell cycle protects KrasG12D cells from cell elimination *in vivo* and *in vitro*. Cell stress responses are major triggers of cell dormancy (*66*). It is possible that non-eliminated KrasG12D cells enter a dormant state in response to competition-induced stress, overriding cell elimination and surviving in tissues. Studies in *Drosophila melanogaster* models demonstrate that cell competition outcomes are a consequence of cellular stress responses activated in ‘loser’ cells (*67–72*). Taking these studies into consideration, our transcriptomic data infer that non-eliminated KrasG12D cells upregulate gene signatures reminiscent of ‘loser’ phenotypes (e.g., Glycolysis (*7*), Unfolded protein response (*73*), p53 (*12, 42, 74–77*), oxidative stress (*78*), NRF2 pathway (*68, 78, 79*)). Thus, cell dormancy may act to restrict expansion of ‘unfit’ KrasG12D cells in the absence of an efficient tissue clearance mechanism and without drastically reducing cell numbers. This would be particularly relevant to adult pancreas, which has limited proliferative capacity (*80*). Indeed, we never observe compensatory proliferation in neighbours following mutant cell clearance (*4*) (Fig. 3B). Future studies are required to determine whether normal-mutant interactions in the adult pancreas induce cellular stress responses in genetically mutant cells and whether this in turn activates cell dormancy. Interestingly, Wnt5a has been shown to induce and maintain cancer cell dormancy in metastatic niches by negatively regulating canonical Wnt/β-catenin signalling (*45, 46*). An exciting future direction of this work will be to elucidate whether the cell dormancy phenotype we describe here is regulated via Wnt-dependent mechanisms described for dormant metastatic cancer cells. Transcriptional profiling of non-eliminated KrasG12D cells support a mechanism whereby Wnt signalling at KrasG12D-normal cell-cell boundaries is not driving cell proliferation. We also show that inhibition of Wnt restores cell extrusion and cell segregation of cell cycle arrested RasV12 cells from normal cells *in vitro*, suggesting a mechanistic link between active Wnt and cell dormancy *in vivo* is plausible.

*TP53* is an important regulator of cell competition. In MDCK cells (*76, 81*) and in developing embryos (*74, 77*), mutant-normal cell competition triggers an elevation in p53 levels and p53 activity in prospective ‘losers’ and this is required for mutant cell elimination (*77*). Studies in the adult hematopoietic system (*82, 83*) and oesophagus (*13, 84*) illustrate that competition outcomes are dependent on relative rather than absolute levels of p53 and are apparent only when cells are under mild stress. Whether cell competition outcomes we describe here are triggered by differentials in p53 activity between mutant and wild type neighbours requires further investigation. Interestingly, we find that co-expression of p53R172H and KrasG12D in the same cell (KPC), provides all mutant cells with a survival advantage and ‘double mutant’ KPC cells are never eliminated. In MDCK cells, ectopic expression of p53R175H/p53R273H mutations in few cells surrounded by normal cells triggers necroptosis and elimination of the p53 mutant cells (*42*); however, expression of p53R273H mutations in RasV12 monolayers prevents p53 mutant cell death (*42*). Recent studies in zebrafish larval skin (*85*) show that while RasG12V-expressing cells are actively eliminated, cells expressing both RasG12V and Tp53R175H mutations are not eliminated and are viable *in vivo*. Both studies support our observations and suggest ‘double mutant’ KPC cells may override cell death signals *in vivo*. Indeed, our RNA sequencing data suggest cells expressing p53R175H mutations downregulate the apoptosis pathway (Fig. 2B, SFig. 2E).

*KRAS* and *TP53* mutations are key drivers of pancreatic ductal adenocarcinoma. Single cell profiling of pancreatic tumorigenesis in mouse models demonstrates that *Kras* mutant cells are very heterogeneous and show signatures of tumorigenesis even before pre-malignant lesions are established (*86*). Our RNA sequencing data confirm this early transformation and cellular heterogeneity profile of KrasG12D cells. Pancreatic cancer is a devastating disease, which is generally diagnosed at late and incurable stages. An improved understanding of the biology of how pancreatic cancer starts and grows in adult tissues will inform the development of new early detection methods. Our results challenge the current understanding of how cancers start in the adult pancreas and suggest that genetically mutant cells must override tissue homoeostasis mechanisms to survive in tissues, prior to transformation/expansion.

## Author Contributions

Conceptualization, CH; Data curation, BS-B; Experimental design, BS-B, CH; Funding acquisition, CH; Investigation & Analysis: BS-B, CH, JPM, OJS; Methodology, BS-B, MA; Validation, BS-B; Writing original draft, BS-B, CH; Writing review & editing, BS-B, CH; Resources, JPM, OJS; Supervision, CH.

## Acknowledgements

We thank Angela Marchbank and colleagues at School of Biosciences Genome Hub (Cardiff University) for technical support with RNA sequencing experiments. We thank Toby Phesse and Valerie Meniel (European Cancer Stem Cell Research Institute, School of Biosciences, Cardiff University) for Wnt antagonists, PCR primers and helpful discussions on the Wnt pathway. We thank F. Afonso (European Cancer Stem Cell Research Institute, School of Biosciences, Cardiff University) for critical reading of the manuscript and helpful discussions. B.S. and M.A. were supported by Cancer Research UK (CRUK) Early Detection project grant (A27838) to C.H. O.J.S. and J.P.M. were supported by Cancer Research UK core funding to the CRUK Scotland Institute (A17196 and A31287) to O.J.S. laboratory (A21139) and to J.P.M. laboratory (A29996). B.S-B. is currently funded by a Pancreatic Cancer UK Career Foundation Fellowship (176937364). This study was supported by Cancer Research UK (CRUK) Early Detection project grant (A27838) to C.H. The manuscript was critically reviewed by Catherine Winchester (CRUK Scotland Institute).

## Methods

### Mouse lines, Induction of Cre recombinase *in vivo*, Inhibition of Wnt *in vivo*

*Pdx1-Cre^ERT^*(*87*)*, LSL-Kras^G12D/+^*(*88*), LSL-*Trp53^R172H/+^* (*89*)*, Rosa26^LSL-tdRFP^* (*90*) mouse lines have all been previously described. Animals were housed in conventional pathogen-free animal facilities and experiments were conducted in accordance with UK Home Office regulations (ASPA 1986 & EU Directive 2010) under the guidelines of Cardiff University Animal Welfare and Ethics Committee. Mice were genotyped by PCR analysis following standard methods and using primer sequences in (*4*). Both male and female 6-8-week-old experimental and control mice were injected with intraperitoneal injection of tamoxifen in corn oil as previously described (*4*). Low recombination levels were obtained with a tamoxifen dose of 1µg/40g bodyweight once, while high recombination (megadose) was induced by three injections of 9mg/40g over 5 days (*4*). Pancreata were harvested at 7-, 35-, 70- or 140-days post tamoxifen induction. To test the functional role of Wnt signalling *in vivo*, 6-8-week-old experimental and control mice were induced with low dose tamoxifen, aged to 35-days post induction of Cre recombinase (p.i.) and treated with WNT-974 1.5mg/kg or vehicle (DMSO) in corn oil by oral gavage, five days per week for four weeks. At the end of the treatment pancreatic tissue was harvested and treated as described below. No statistical method was used to pre-determine sample size. For most animal studies, experiments were not randomized, and investigators were not blinded to allocation during experiments. To test the functional role of Wnt signalling mice were randomly allocated into vehicle or WNT-974 treated groups. All experiments were reproduced using at least three animals of each genotype.

### Tissue staining

For histological observation, pancreas was harvested at specified timepoints and fixed in 10% neutral buffered formalin overnight at 4°C. Then pancreas was dissected into two pieces (head-body/body-tail) and embedded either in paraffin or OCT matrix. FFPE embedded pancreas was sectioned (7µm thickness) and stained with specific antibodies (table 1). Anti-RFP, anti-cleaved caspase-3, and anti-Ki-67 staining were performed via immunohistochemistry (IHC) in FFPE tissue sections. Tissues sections were dewaxed and rehydrated. For antigen retrieval, tissue sections were incubated for 15 minutes at 37°C in 20ug/ml Proteinase K diluted in TBS-T or boiled in citrate buffer pH 6 for 15 minutes (specified in table 1). Tissues were then blocked with 3% H_2_O_2_ for 20 or 10 minutes at room temperature (depending on the antigen retrieval method) respectively and 5% NGS TBS-T for 30-60 minutes at room temperature. Sections were then incubated overnight at 4°C with the primary antibody diluted in 5% NGS TBS-T (see dilutions in table 1). Tissues were stained with the secondary antibody ImmPRESS goat anti-Rabbit (MP-7451, Vector Labs) for 60 minutes at room temperature, followed by DAB chromogen (Peroxydase substrate kit, Vector Labs). Sections were dehydrated and mounted with DPX (Sigma-Aldrich) and scanned using the Axio Scan Z1 slide scanner (Zeiss) 20X magnification. Immunofluorescence (IF) co-staining using either anti-CD44 and anti-RFP (Rockland) was performed in FFPE tissues slices using the same antigen retrieval and primary antibody incubations as described for IHC, followed by Alexa fluor secondary antibody incubation for 60 minutes at room temperature and Hoechst (ThermoFisher) (table 1) and mounted using Mowiol (Sigma-Aldrich).

**Table 1.**

| Antibody and fluorescence stains | Application | Antigen Retrieval | Dilution | Source | Identifier |
| --- | --- | --- | --- | --- | --- |
| Ki-67 | IHC, IF | ProK 15' | 1:500 | Abcam | ab16667 |
| CC-3 | IHC | Citrate pH6 15' | 1:300 | Cell Signaling | 9661s |
| RFP | IHC, IF | ProK 15' | 1:500 | Rockland | 600-401-379 |
| Lectin PNA-A488 | FACS |  |  | Invitrogen | L21409 |
| c-Myc | WB |  | 1:500 | Santa Cruz | sc788 |
| Phospho-p38 | WB |  | 1:1000 | Cell Signaling | 9211s |
| p38a | WB |  | 1:1000 | Santa Cruz | sc136210 |
| Vinculin | WB |  | 1:1000 | Sigma Aldrich | V4505 |
| Phosphor-ERK | WB |  | 1:1000 |  |  |
| ERK | WB |  | 1:1000 | Cell Signaling |  |
| p21WAP/Cip | WB |  | 1:1000 | Sigma Aldrich | P1484 |
| CD44 | IF | ProK 15' | 1:25 | Cell Signaling | 3570S |
| K-cadherin | IF | PoK 15' | 1:50 | ThermoFisher | MA1-06305 |
| β-catenin | IF | Citrate pH6 15' | 1:50 | BD Biosciences | 610154 |
| RFP | IF | Citrate pH6 15' | 1:500 | Creative Diagnosis | DPATB-H83194 |
| E-cadherin | IF | Citrate pH6 15' | 1:1000 | BD Biosciences | 610182 |
| p27 | IF |  | 1:25 | Santa Cruz | SC-528 |
| Wnt5a | IF |  | 1:50 | Santa Cruz | SC-30224 |
| Dvl2 | IF |  |  | Santa Cruz | SC-13974 |
| goat anti-rabbit AF568 | IF |  | 1:1000 | ThermoFisher | A-11011 |
| donkey anti-mouse AF488 |  |  | 1:1000 | ThermoFisher | A-21202 |
| goat anti-rabbit AF488 | IF |  | 1:1000 | ThermoFisher | A-11008 |
| goat anti-rabbit HRP | WB |  | 1:5000 | Vector Laboratories | MP-7451-15 |
| goat anti-mouse HRP | WB |  | 1:5000 | Vector Laboratories | MP-7802-15 |
| Phalloidin | IF |  | 1:1000 | Sigma Aldrich | 65906 |
| E-cadherin | F |  | 1:100 | ThermoFisher | 13-1900 |
| Caveolin | IF |  | 1:50 | Abcam | Ad2910 |
| Hoechst |  |  | 1:1000 | ThermoFisher | H3570 |

For β-catenin E-cadherin, anti-K-cadherin, and RFP (Creative Diagnostics) IF staining, pancreas was embedded in butyl-methyl methacrylate plastic (BMMA) under UV following dehydration and resin infiltration (*4*). Tissue was sectioned in 2µm thick slices and rehydrated. Tissue staining followed the same protocol as IF in FFPE sections. Antigen retrieval and antibody concentrations are listed in table 1.

Pancreas embedded in OCT matrix was sectioned (10µm thickness), permeabilized with 0.05% SDS (Sigma-Aldrich) and 0.05% Triton X100 (Sigma-Aldrich) solution and stained with anti-p27, anti-Wnt5a, anti-Dvl2 primary antibodies (table 1) for 2 hours at room temperature, Alexa fluor secondary antibody incubation for 60 minutes at room temperature and Hoechst (ThermoFisher) (table 1), and mounted using Mowiol (Sigma-Aldrich).

Global levels of endogenous RFP were measured in 10µm formalin fixed OCT frozen sections, as previously described (*4*). RFP positive area was averaged from 3 tissue slices per mouse separated by at least 20µm between each section. Fluorescence imaging was carried out on a Zeiss LSM710 confocal microscope.

To assess the presence of PanIN lesions, FFPE sections were stained with Alcian blue as previously described (*4*). β-galactosidase activity was determined using the Cell Signaling Senescence β-galactosidase Staining kit (9860) using the manufacturers’ instructions.

### Image analysis

The percentage of RFP+ tissue area was determined using endogenous RFP levels quantified as described in (*4*). Using Fiji (RRID:SCR_002285) software,(*91*) positive areas were thresholded using RFP fluorescence. Tissue autofluorescence was used to determine total tissue area. The percentage of RFP+ ducts were analysed using E-cadherin/RFP/Hoechst-stained sections to identify ducts. At least 20 ducts were quantified per mouse and ducts were considered positive when they contained at least one RFP+ cell. The percentage of RFP+ cells in pancreatic islets were scored in FFPE IHC sections stained with anti-RFP antibody and using ‘positive cell count’ tool in QuPath-0.4.1 software. RFP positive cluster size, perimeter and area were determined in FFPE IHC sections stained with anti-RFP antibody and using QuPath-0.4.1 software by analysing shape features of RFP positive (DAB positive) areas. Ki-67 positive and Cleaved Caspase-3 positive cells were analysed using QuPath-0.4.1 positive cell count feature.

Wnt5a fluorescence intensity in the tissue was quantified by masking the tissue area (AF568) and measuring mean fluorescence intensity for Wnt5a (AF488). CD44, β-catenin and E-cadherin fluorescence intensity at the border between RFP positive and negative cells were quantified by measuring mean fluorescence intensity of 2.43×2.43-pixel circles (ROIs) along the membrane (RFP+/RFP-) separated by approximately 5 pixels. CD44, Dvl2 and K-cadherin fluorescence in clusters was quantified by masking RFP+ (AF568) areas and quantifying mean fluorescence intensity for CD44/Dvl2/K-cadherin (AF488). β-catenin fluorescence in nuclei was quantified by masking nuclei (Hoechst) and quantifying mean fluorescence intensity for β-catenin (AF488) in RFP+ cells. To avoid noise associated with tissue autofluorescence, fluorescence intensity measurements were made relative to image global mean fluorescence intensity (AF488).

### Pancreas digestion and RNA Sequencing

Pancreas tissues were harvested 35 days post low-dose tamoxifen induction. To reach the minimum of 5000 cells and a minimum of 100ng RNA/sample for bulk RNA sequencing, we combined three pancreas tissues of the same genotype to produce one sample per genotype. Each sample was replicated three times to generate three biological repeats per genotype for sequencing. Pancreas tissues were digested as previously described (*92*). Using an EGTA-based buffer the pancreas from each mouse was inflated and slightly digested with collagenase. The tissue was chopped and further digested using a calcium-based buffer with collagenase. Single cell digested tissue was stained with lectin PNA-A488 (ThermoFisher) and DAPI (ThermoFisher Scientific). DAPI negative, lectin positive, RFP positive cells were sorted using the FACS Aria Fusion sorter. A minimum of 5,000 cells were sorted for each sample. RNA was extracted using the RNeasy Micro kit (Qiagen) following the manufacturer’s recommendations. Paired-end sequencing was performed using Sanger sequencing Illumina 1.9. Sequenced reads Fastq files were quality-checked using FastQC. Reads were aligned to the mouse genome (Ensembl-GRCm38 *Mus musculus*) using the STAR package, and reads were counted using the FeatureCounts package, after removing duplicates using MarkDuplicates from GATK. Differential expression was normalised and calculated using Deseq2, comparing the different genotypes against the Control. Finally, GSEA 4.2.2 software was used for enrichment analysis of different pathways and GraphPad Prism 10.0.2 for Heatmap graphs and normalised enrichment score (NES) graphs. The RNAseq data generated during this study are available at GEO ID: XX.

### MDCK cell lines and *In vitro* experiments

To assess the interactions between RasV12 and MDCK cells, MDCK-pTR GFP–RasV12 cells were combined with parental MDCK cells at a ratio of 1:50. Treatment with tetracycline (2µg/mL) induces GFP-RasV12 expression in MDCK-pTR GFP–RasV12 cells (*5, 9*). Cells were plated at a density of 2.5 x 10^5^ cells per well in MW24 plates (Corning) carrying 18mm diameter glass cover slips (VWR). Mixed cells were incubated for 30h or 48h at 37°C before fixed with 4% Formaldehyde (ThermoFisher Scientific). Cells were then permeabilized and stained (table 1). GFP-RasV12 cells were considered non-extruded when more than 30% of their cytoplasm was basally protruded. Three technical replicates per experiment in three experiments (i.e., using three independent frozen stocks of parental and GFP-RasV12 MDCK cells) were performed and at least 150 GFP-RasV12 cells were quantified per replicate.

For c-Myc silencing, tetracycline induced GFP-RasV12 cells were transfected with 100ng siRNA oligos targeting *Myc* using Lipofectamine 3000 (Invitrogen) (siRNA siMyc1: AAGACGUUGUGUGUUCGCCUC – as GAGGCGAACACACAACGUCUU, siRNA siMyc2: AAUUUCAACUGUUCUCGCCGC – as GCGGCGAGAACAGUUGAAAUU and siScr)^24^. For cell extrusion and proliferation experiments, GFP-RasV12 cells were trypsinized and mixed with parental MDCK cells 24h after siMyc1+2 transfection and were then fixed following a further 48h.

For experiments comparing PBS, Wnt3a and Wnt5a treatments effect, RasV12 cells were mixed 1:50 with parental MDCK cells and once cells were set, induced with tetracycline. Two hours post-tetracycline induction cells were treated with 1µg/mL of human recombinant Wnt3a (Peprotech), 100ng/mL of human recombinant human/mouse Wnt5a (R&D Systems) or PBS (control-Sigma-Aldrich). Cells were fixed 30h after PBS, Wnt3a or Wnt5a treatment for cell extrusion analysis and for analysis of E-cadherin/β-catenin at the RasV12-MDCK contact or within GFP-RasV12 cell clusters. E-cadherin endocytosis and Caveolin fluorescence were assessed by fixing cells 16h after PBS/Wnt5a treatment.

For Wnt pathway inhibition experiments, tetracycline induced GFP-RasV12 cells were transfected with siMyc1+2/siScr or treated with recombinant Wnt5a/PBS for 24h and then mixed with parental MDCK cells. Wnt5a treatment was maintained during the whole experiment. WNT-974 (Stratech) porcupine inhibitor (1µM) or *OMP*-18R5 (10mg/mL) was added to the medium 8h after GFP-RasV12 and MDCK cells were mixed and plated. DMSO treated GFP-RasV12 cells were used as control for both inhibitors. To assess cell extrusion, cells were fixed 48h after plating. E-cadherin endocytosis was assessed by mixing RasV12 cells with MDCK parental cells, then RasV12 expression was induced by tetracycline and treated with PBS or Wnt5a 2h after. Four hours after PBS/Wnt5a treatment, cells were treated with DMSO or WNT-974. Cells were fixed 16h after PBS/Wnt5a treatment.

Immunofluorescent staining was performed by permeabilization of cells with 0.05% SDS (Sigma-Aldrich) and 0.05% Triton X100 (Sigma-Aldrich) solution for 15 minutes. Cells were stained with anti-Ki-67, anti-E-cadherin, anti-β-catenin or Caveolin primary antibodies for 2h at room temperature, Alexa fluor secondary antibody incubation for 60 minutes at room temperature and Hoechst (ThermoFisher) (table 1) and mounted using Mowiol (Sigma-Aldrich). For proliferation analysis at least 30 extruded and 30 non-extruded cells were quantified in siMyc 1+2 treated cells per experimental replicate. For E-cadherin, β-catenin or Caveolin at least 10 GFP-RasV12 cell-clusters per experimental replicate were quantified. Fluorescence intensity at the border between RasV12 and parental MDCK cells were quantified by measuring mean fluorescence intensity of 2.27×2.27-pixel circles (ROIs) along the membrane separated by approximately 5 pixels. E-cadherin endocytosis was measured quantifying the mean fluorescence intensity of the cytoplasm of each RasV12 cell. At least 10 cells were quantified in three technical replicates per experiment in three experiments.

Confrontation assays and migration speed analysis were carried out as described (*9*). 1×10^4^ cells were plated in inserts (Ibidi) in MW24 plates (Corning), allowed to form monolayers (8h) before removing the insert and moving the plate to IncuCyte S3 (Sartorius) for live cell imaging. Images were captured every 15min for 48h. siMyc/siScr-GFP-RasV12 cells were plated 24h after transfection and inserts were removed 8h post transfection. For Wnt5a experiments cells were treated with PBS/Wnt5a for 8h before inserts were removed. PBS/Wnt5a treatment was maintained during the whole experiment. WNT-974 porcupine inhibitor (Stratech)/DMSO (Sigma-Aldrich) was added to the medium when inserts were removed. Retraction was measured as the distance GFP-RasV12 cells migrated during 24h, following initial cell-cell collision with MDCK cells. Coefficient of boundary smoothness was measured using Fiji in cell confrontation assays at the end of the 24 h experiment and as previously described (*9*). We performed three technical replicates per experiment in three experiments.

Western blotting was performed using GFP-RasV12 cell lysates transfected with siMyc/siScr siRNAs at 48h post transfection (SFig. 3E), treated for 48h with WNT-974/DMSO 24h post-transfection (SFig. 5D). Cells were lysed using laemmli buffer (0.0625 M Tris base, 2% SDS, 10% glycerol). Proteins were separated on 12% polyacrylamide gels under reducing PAGE conditions. Proteins were then transferred to PVDF membranes (Immobilon-P, Merck) using wet transfer (1h 100V). Membranes were blocked in 5% milk TBS-T for 1h and antibodies were added in the same buffer at the concentrations listed in table 1 for overnight at 4°C or 2h at room temperature. Secondary antibodies (table 1) were added in 5% milk TBS-T and membranes were developed using ECL luminol kit (Merck) and chemiluminescence films (Amersham Hyperfilm ECL, GE Healthcare). Protein levels were quantified using Fiji (RRID:SCR_002285) software ‘Gels’ tool.

Real time cell confluency was measured using Incucyte S3 live cell imaging instrument and software (v2020). 1×10^3^ cells siMyc/siScr-GFP-RasV12 cells were plated 24h after transfection with siMyc/siScr siRNAs in MW96 (Corning). Images were captured every 15min for 48h. We performed three technical replicates per experiment in three experiments.

### Statistical tests

Statistical analyses were performed using GraphPad Prism 10 software. Normally distributed data, as determined by the Shapiro-Wilke test or D’Agostino and Pearson test were analysed using unpaired Student’s t tests. A *p* value of <0.05 was taken as significant and a rejection of the null hypothesis. No statistical method was used to pre-determine sample size. Definition of *n* is in the figure legends. Gene set enrichment analysis (GSEA) was performed using GSEA 4.2.2 software. GSEA was used to identify gene sets that were differentially expressed between two groups of samples. Gene sets were considered enriched if they had a false discovery rate (FDR) of less than 0.25. Heatmaps were created by computing normalised gene counts for each individual sample into row z-scores.

---

**Supplementary figure 1.**
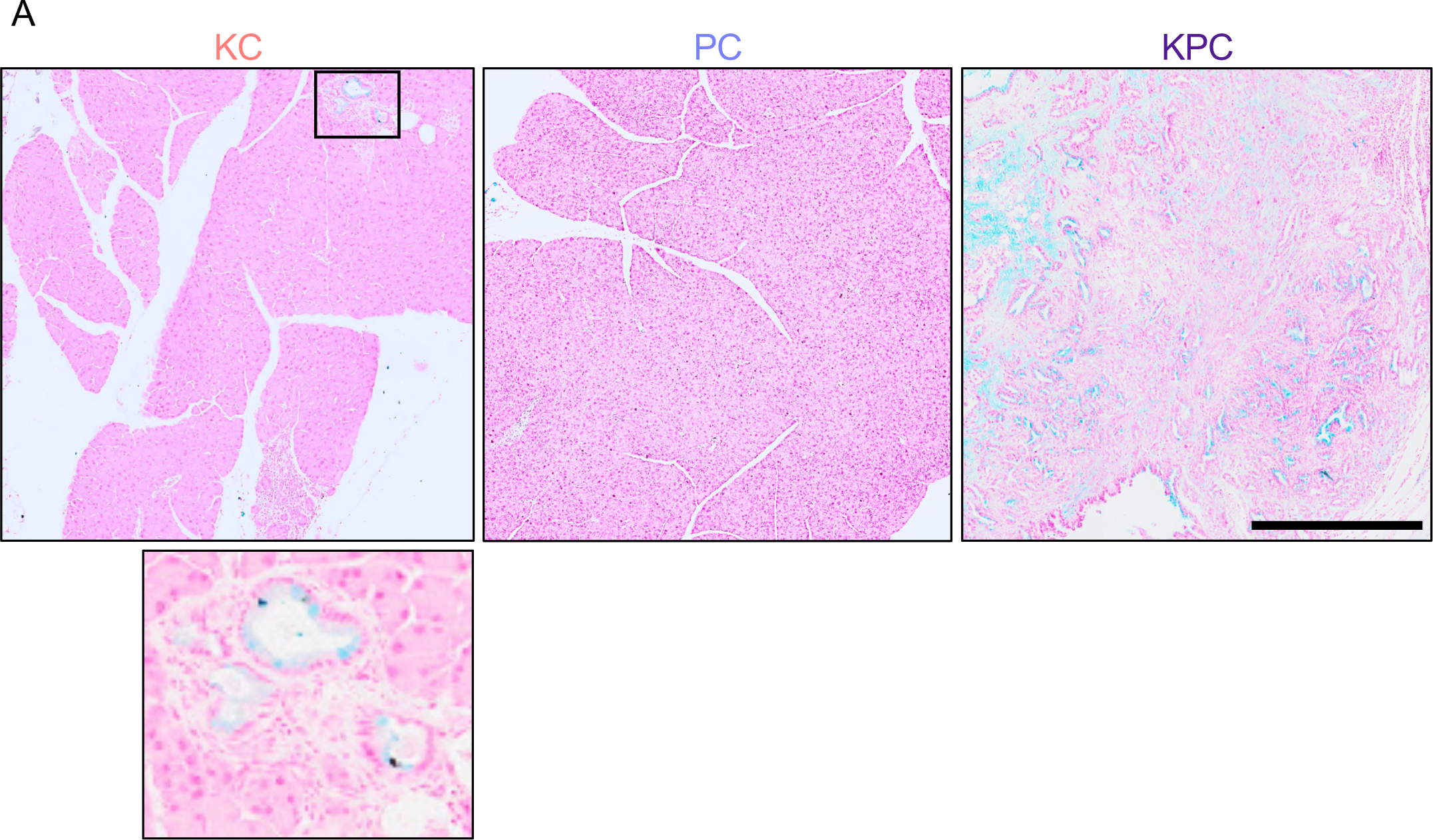
Non-eliminated mutant cells can progress to mucin positive lesions. **A** Pancreas tissues harvested from KrasG12D (KC), p53R172H (PC) or double mutant (KPC) expressing mice at 168 days p.i. and stained for mucin (using Alcian blue). Scale bar, 500 µm. Boxed area shown magnified (lower panel).

**Supplementary figure 2.**
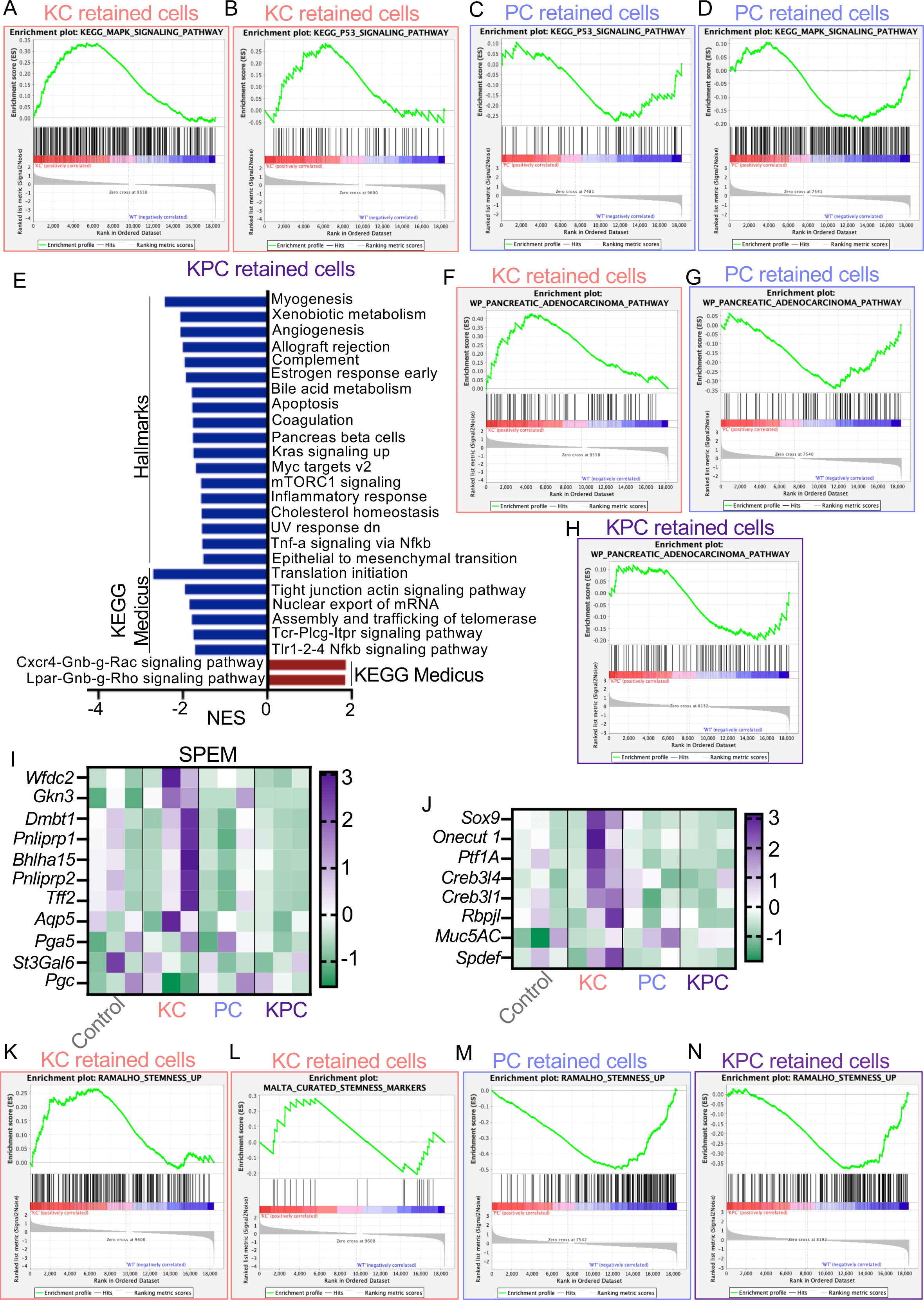
Transcriptomic analysis indicates Kras and p53 pathways deregulation in retained cells. Non-eliminated KrasG12D (KC) are stem-like, whereas p53R172H-expressing populations are not. GSEA enrichment plots showing **A** KEGG_MAPK_signaling_pathway (M10792) and **B** KEGG_p53_signaling_pathway (M6370) positive correlation in KrasG12D (KC) cells compared to Control (WT) and **C** KEGG_p53_signaling_pathway (M6370) and **D** KEGG_MAPK_signaling_pathway (M10792) negative correlation in p53R172H (PC) cells compared to Control (WT). **E** Normalised enrichment scores (NES) of gene set enrichment analysis (GSEA) on the Hallmark and KEGG Medicus gene sets for the RNA sequencing experiment analysis of KPC retained cells. Only gene sets with an FDR of less than 0.25 were included in the graph. The complete list that contains the results of GSEA analysis is provided in Supplementary Tables S8-S9. GSEA enrichment plots of WP_pancreatic_adenocarcinoma_pathway (M39732) positively correlated in **F** KrasG12D (KC) cells, and negatively correlated in **G** p53R172H (PC) cells and **H** KrasG12D p53R172H (KPC) cells compared to Control (WT). **I-J** Heatmaps showing gene expression in non-eliminated KrasG12D (KC) cells and control of **I** spasmolytic polypeptide-expressing metaplasia and **J** pancreatic lineage. Heatmaps show row z-scores for the expression of each gene for three samples (three pooled mice per sample) per genotype obtained from the RNA sequencing experiment. Genes are listed in rows, genotypes in columns. GSEA enrichment plots indicate positive correlation in KrasG12D (KC) retained cells compared to Control (WT) for **K** Enrichment of Ramalho_stemness_up (M9473) and **L** Malta_curated_stemness_markers (M30411) (embryonic, neural, and hematopoietic stem cells). GSEA enrichment plots indicate negative correlation for Ramalho_Stemness_up (M30411) in **M** p53R172H (PC) retained cells and **N** KrasG12D p53R172H (KPC) retained cells compared to Control (WT).

**Supplementary figure 3.**
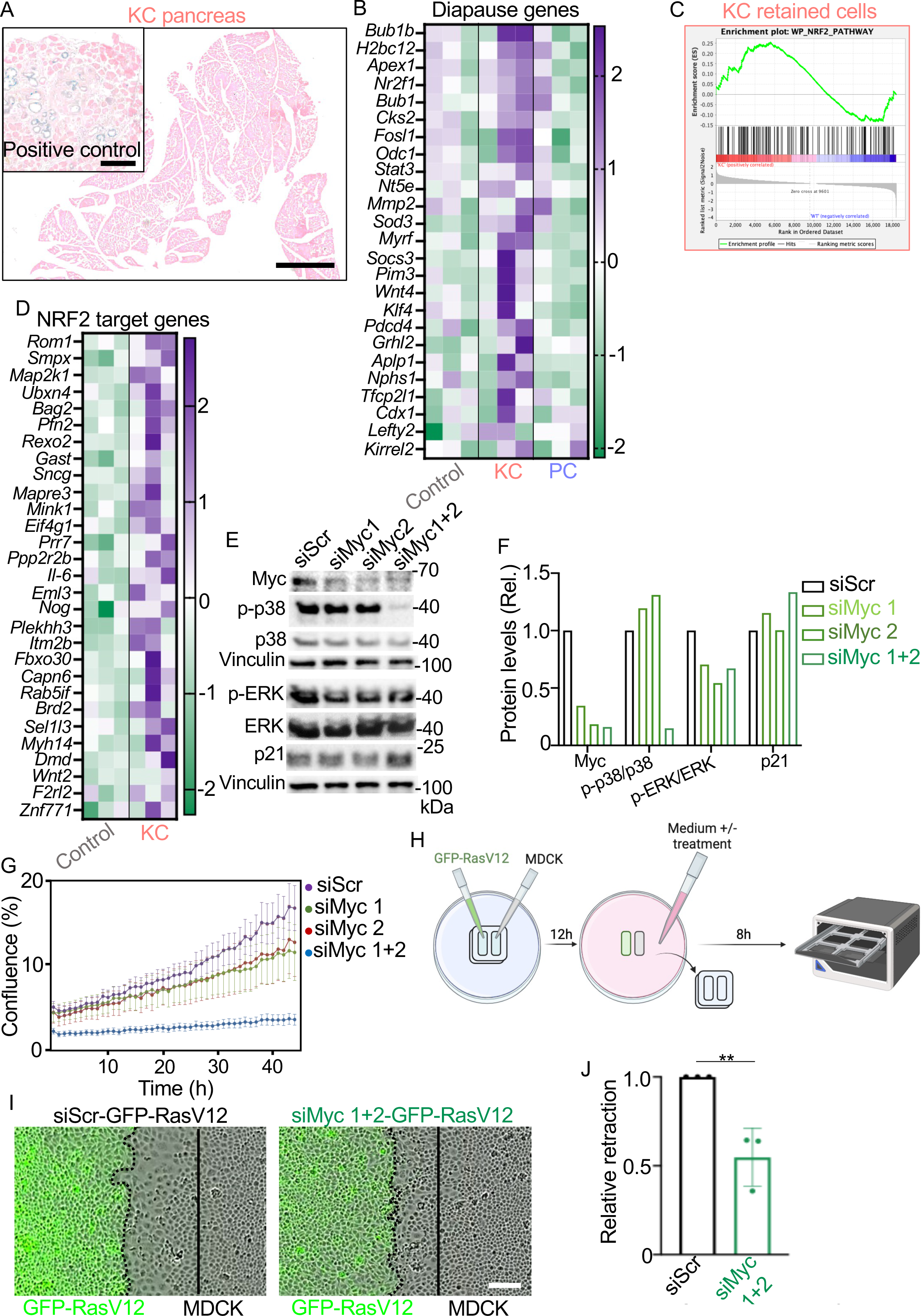
Non-eliminated KrasG12D cells upregulate cell dormancy-associated pathways. Cell cycle arrest abrogates RasV12 cell extrusion *in vitro*. **A** Representative images of β-galactosidase staining of pancreas tissue harvested from a KrasG12D (KC) mouse at 35 days p.i (scale bar, 1000µm) and in PanIN lesions as positive control (scale bar, 250 µm). **B** Diapause gene expression (rows) in Control, KrasG12D (KC) and p53R172H (PC) cell transcriptomes (columns). The heatmap shows row z-scores for each gene obtained from three samples per genotype (three pooled mice per samples) from the RNA sequencing experiment. **C** GSEA enrichment plots showing positive correlation in KrasG12D (KC) retained cells compared with Control (WT) for WP_NRF2_pathway (M39454). **D** Heatmap of expression levels of NRF2 target genes (rows) in three KrasG12D (KC) and three Control samples (three pooled mice per sample). The heatmap shows row z-scores for each gene obtained from the RNA sequencing experiment. Only genes with z-scores of a foldchange of 2.25 are shown. The full list of genes can be found in supplementary table S12. **E** Immunodetection of indicated antigens in siScr-GFP-RasV12 (siScr), siMyc1-GFP-RasV12 (siMyc1), siMyc2-GFP-RasV12 (siMyc2) and siMyc1+2-GFP-RasV12 (siMyc1+2) cells protein lysates. GFP-RasV12 cells were first transfected with scrambled siRNA (siScr) or two siRNA oligos targeting endogenous *Myc* (siMyc1, siMyc2 or combined siMyc1+2). Lysates were collected 48h after transfection. Two blots were run, and vinculin was used as loading control for each blot (vinculin band shown at the bottom of each blot). p-p38=phospho-p38; p-ERK=phospho-ERK. **F** Quantification of protein levels in the blot in E. Values are relative to siScr protein levels. **G** Real time cell confluence of GFP-RasV12 cells transfected with different siRNAs as quantified via Incucyte S3 imaging. Cell confluence was determined in cells 12h post siRNA transfection over 48h. Data indicate mean and +/- s.d. for three experiments per condition. **H** Schematic representation of cell confrontation assay experiments. Illustration created with BioRender.com. **I** Representative time-lapse images of cell confrontation assays. GFP-RasV12 cells expressing either scramble siRNA (siScr-GFP-RasV12) or combined Myc1+2 siRNA (siMyc-GFP-RasV12) confront non-labelled parental MDCK cells. Dashed lines highlight the border between GFP-RasV12 cells and MDCK cells. Solid lines indicate MDCK cells’ migration front at the beginning of the experiment. Scale bar, 100µm. **J** Relative retraction distance of GFP-RasV12 cells transfected with either scrambled siRNA (siScr-GFP-RasV12; black bar) or combined Myc1+2 siRNA (siMyc-GFP-RasV12; green bar) following collision with parental MDCK cells until 24h later. Results are presented as retraction distance in siMyc-GFP-RasV12 relative to siScr-GFP-RasV12. Data represent mean and +/- s.d. measurements of three experiments. Student’s t test was used to analyse the data, **p<0.005.

**Supplementary figure 4.**
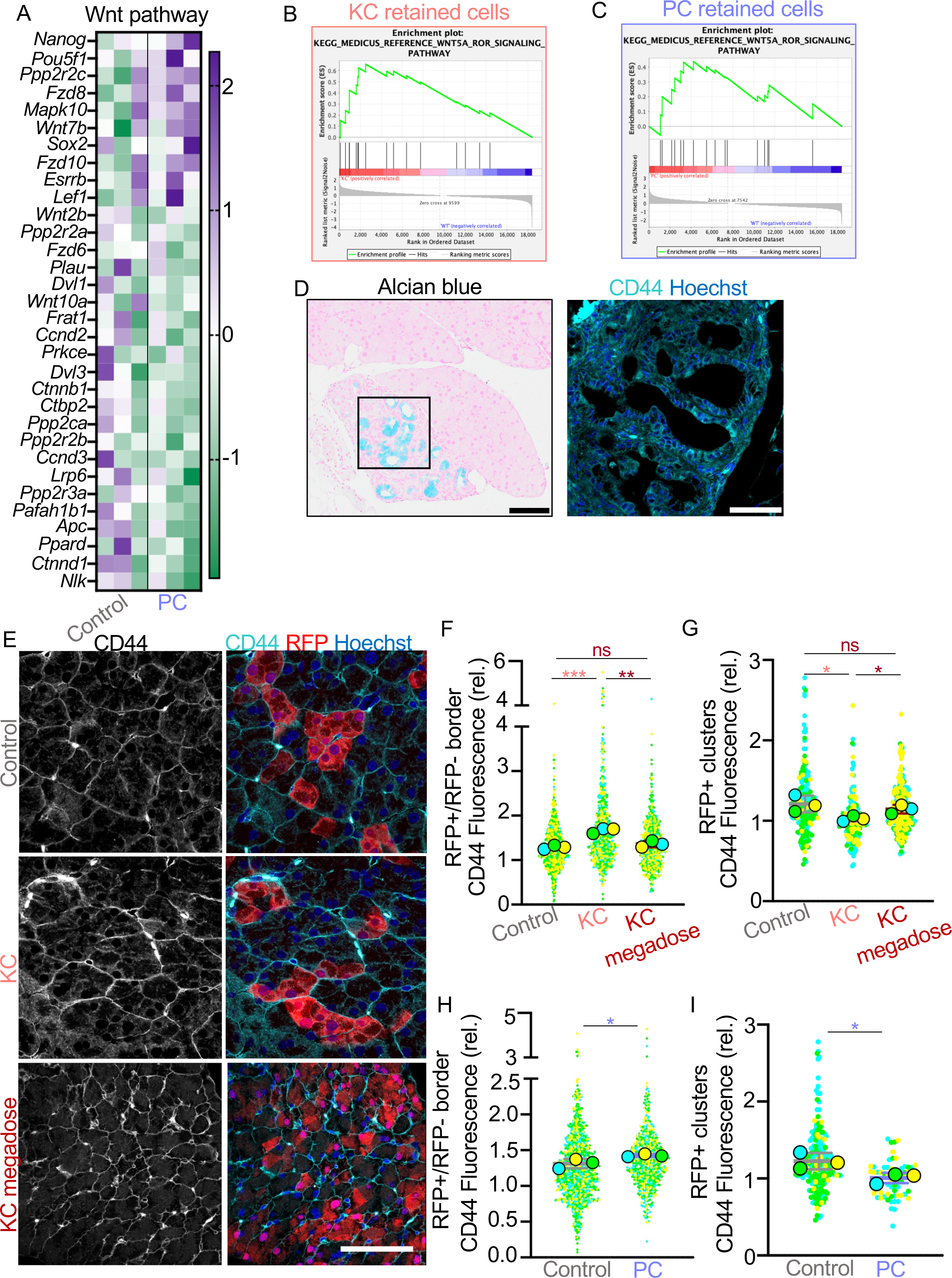
Wnt pathway is active in non-eliminated KrasG12D or p53R172H cells. **A** Heatmap of Wnt signalling pathway related genes (rows) in Control and p53R712H (PC) cell transcriptomes (columns). The heatmap shows row z-scores for each gene obtained from the RNA sequencing experiment. Data from three samples (three mice pooled per sample) per genotype are shown. Only genes with z-scores of a foldchange of 2 are shown in the Wnt pathway panel. The full list of genes can be found in supplementary table S11. GSEA enrichment analysis plots showing positive correlation in **B** KrasG12D (KC) and **C** p53R172H (PC) retained cells compared with Control (WT) for the KEGG_MEDICUS_reference_WNT5A_ROR_signaling_pathway (M47822). **D** Representative images of PanINs in pancreas tissue sections harvested from KrasG12D-expressing (KC) mice at 140 days p.i. Left: Mucin positive PanINs (blue). Scale bar, 100 µm. Right: confocal image of anti-CD44 antibody (cyan) labelling and Hoechst (blue) in PanINs in a consecutive tissue slice. Scale bar, 50 µm. **E** Confocal images of pancreas tissues harvested at 35-days p.i. from Control, KrasG12D (KC) and KrasG12D high dose tamoxifen (KC megadose) mice and stained with anti-RFP (red) and anti-CD44 (cyan) antibodies and Hoechst (blue). Scale bar, 50µm. **F** Mean fluorescence intensity of CD44 at the boundary between RFP positive and RFP negative cells in Control, KrasG12D low dose tamoxifen (KC) and KrasG12D high dose tamoxifen (KC Megadose) tissues harvested at 35 days p.i. CD44 fluorescence was quantified and reported as in figure 4I. Fluorescence was measured in ROIs, separated by a constant distance, along the cell-cell boundary. SuperPlot shows all the quantified ROIs (smaller circles) and the mean for each mouse (n=3 samples; larger circles - blue, green, and yellow). The graph shows mean and +/- s.d. for the three mice. Student’s t test was used to analyse the data. **p=0.0047; ***p=0.0009; ns=0.2136. **G** Mean fluorescence intensity of CD44 in entire RFP positive clusters. Mean fluorescence intensity is relative to the background. Fluorescence was measured in ROIs, separated by a constant distance, along the cell-cell boundary. SuperPlot shows all the quantified RFP+ clusters (smaller circles) and the mean for each mouse (n=3 samples; larger circles - blue, green, and yellow). The graph shows mean and +/- s.d. for the three mice. Student’s t test was used to analyse the data. *p<0.05; ns=0.3916. **H** Mean fluorescence intensity of CD44 at the boundary between RFP positive and RFP negative cells in Control and p53R172H (PC) tissues harvested at 35 days p.i. CD44 fluorescence was quantified and reported as in figure 4I. Control quantification is the same used in figure 4I. Fluorescence was measured in ROIs, separated by a constant distance, along the cell-cell boundary. SuperPlot shows all the quantified ROIs (smaller circles) and the mean for each mouse (n=3 samples; larger circles - blue, green, and yellow). The graph shows mean and +/- s.d. for the three mice. Student’s t test was used to analyse the data. *p=0.0470. **I** Mean fluorescence intensity of CD44 in entire RFP positive clusters CD44 fluorescence was quantified and reported as in G. SuperPlot shows all the quantified clusters (circles) and the mean for each mouse (n=3 samples; triangle, square and diamond). The graph shows mean and +/- s.d. for the three mice. Student’s t test was used to analyse the data. *p=0.0397.

**Supplementary figure 5.**
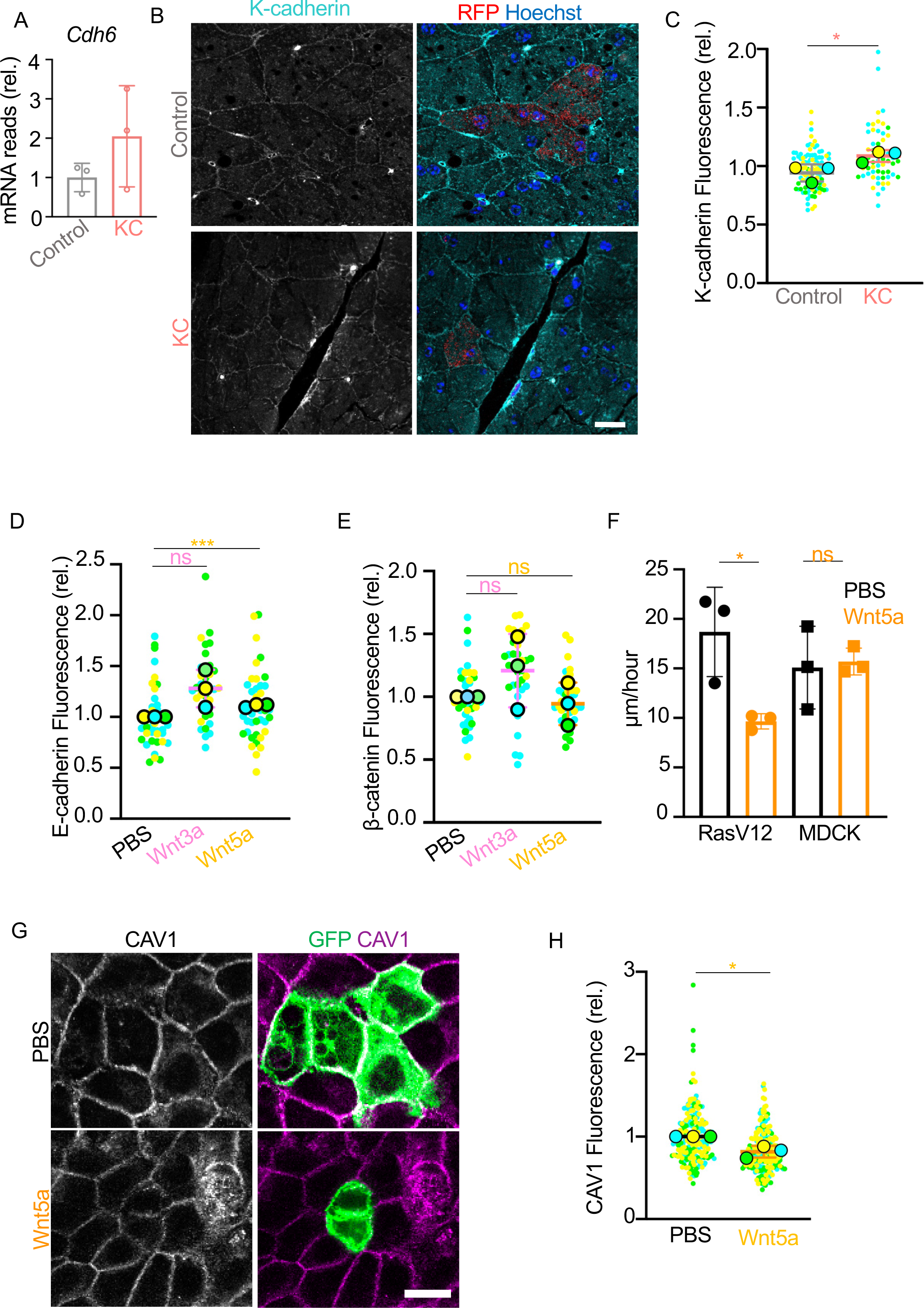
Wnt5a increases RasV12 cell cohesion. **A** *Cdh6* mRNA reads relative to control reads in the three samples in Control and three samples in KrasG12D (KC) obtained from the RNA sequencing experiment. Graph represents mean +/- s.d. reads. **B** Confocal images of pancreas tissues harvested at 35-days p.i. from Control and KrasG12D (KC) mice. Tissues were stained with anti-RFP (red) and anti-K-cadherin (cyan) antibodies and Hoechst (blue). Scale bar, 50µm. **C** Mean fluorescence intensity of K-cadherin clusters in Control or KrasG12D (KC) pancreas tissues harvested at 35 days p.i. SuperPlot shows all the quantified RFP+ clusters (smaller circles) and the mean for each mouse (n=3 samples; larger circles - blue, green, and yellow). The graph shows mean and +/- s.d. for the three mice Student’s t test was used to analyse the data, \**p*=0.0474. **D, E** Mean fluorescence intensity of E-cadherin (**D**) or β-catenin (**E**) in GFP-RasV12 cell clusters (mixed with MDCK cells) treated with PBS (black), Wnt3a (pink) or Wnt5a (yellow) for 30h. SuperPlot shows all the quantified GFP-RasV12 clusters (smaller circles) and the mean for each experiment (n=3 samples; larger circles - blue, green, and yellow). The graph shows mean and +/- s.d. for the three experiments. Student’s t test was used to analyse the data. ***p=0.0003; ns p>0.05. **F** Cell speed (μm per hour) at which RasV12 (circles) or MDCK (squares) cells treated with PBS (black) or Wnt5a (yellow) migrate. The data represent mean +/- s.d. from three independent experiment. Student’s t test was used to analyse the data. *p=0.0268; ns p=0.8202. **G** Confocal images of GFP-RasV12 cells mixed with MDCK cells treated with PBS or Wnt5a for 16h. Cells were stained with anti-CAV1 (magenta). Scale bar, 20µm. **H** Mean fluorescence intensity of CAV1 at the boundary between GFP-RasV12 and MDCK cells treated with PBS (black) or Wnt5a (yellow) for 16h. Fluorescence was measured in ROIs, separated by a constant distance, along the cell-cell boundary. SuperPlot shows all the quantified ROIs (smaller circles) and the mean for each experiment (n=3 samples; larger circles - blue, green, and yellow). The graph shows mean and +/- s.d. for the three experiments. Student’s t test was used to analyse the data. *p=0.0117.

**Supplementary figure 6.**
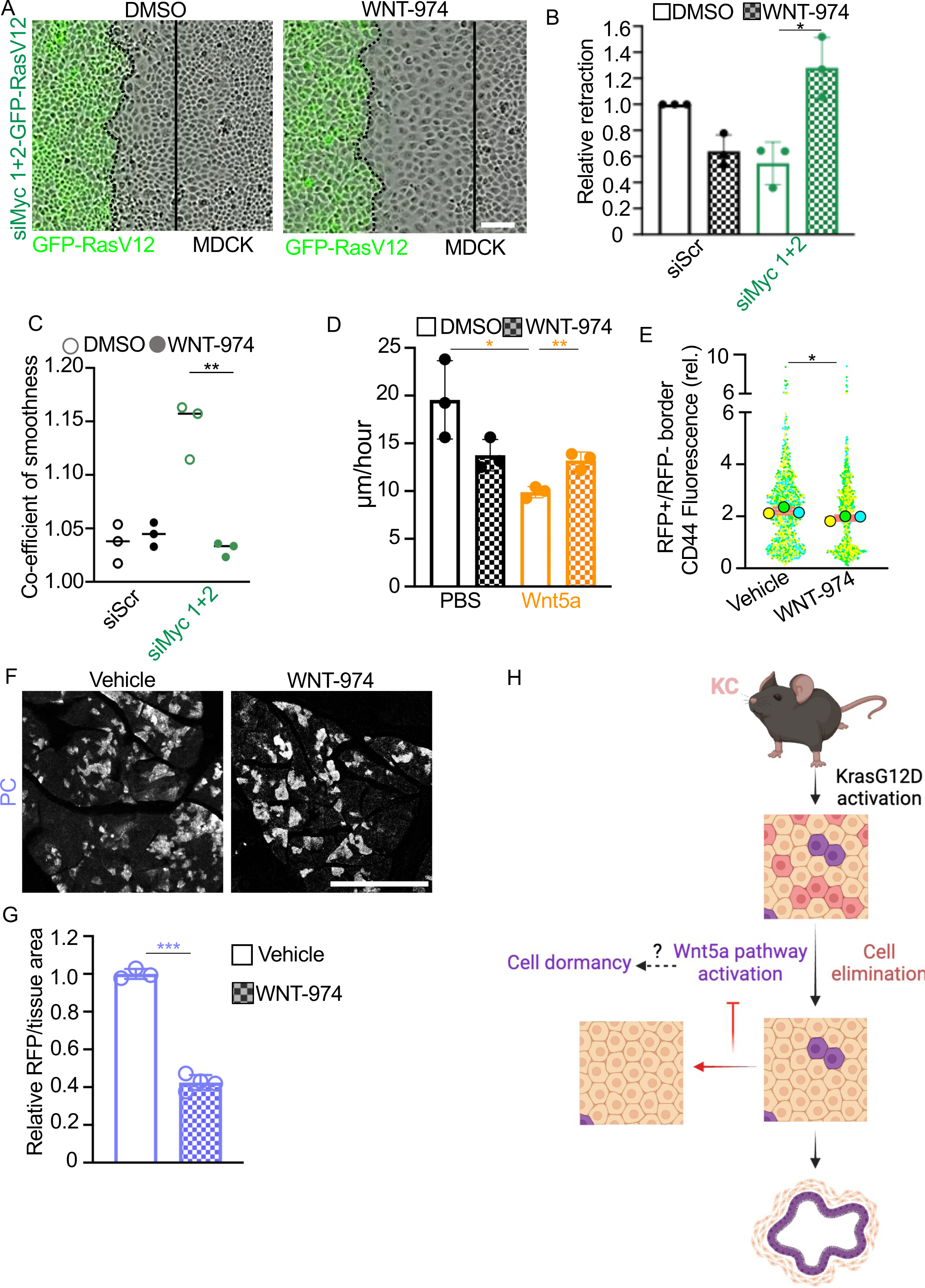
Cell cycle arrest and Wnt signalling prevent GFP-RasV12 segregation from normal MDCK cells *in vitro* and pancreas cell elimination *in vivo*. **A** Representative time-lapse images of cell confrontation assays. GFP-RasV12 cells expressing combined Myc1+2 siRNA (siMyc 1+2-GFP-RasV12) confront non-labelled parental MDCK cells. Assays were conducted in the presence of DMSO or WNT-974. Dashed lines highlight the border between GFP-RasV12 cells and MDCK cells. Solid lines indicate migration front of MDCK cells at the beginning of the experiment. Scale bar, 100µm. **B** Relative retraction distance of GFP-RasV12 cells transfected with scrambled siRNA (siScr; black bars) or two siRNA oligos targeting endogenous *Myc* (combined siMyc1+2: siMyc-GFP-RasV12; green bars) following collision with parental MDCK cells until 24h later. Assays were conducted in the presence of DMSO (solid bars) or WNT-974 (hatched bars). Results are presented as retraction distance in siMyc-GFP-RasV12 relative to siScr-GFP-RasV12. Data represent mean and +/- s.d. of three experiments. Student’s t test was used to analyse the data, *p<0.05. **C** Co-efficient of boundary smoothness in cell confrontation assay. Co-efficient was calculated by measuring the length of the cell-cell boundary between GFP-RasV12 and MDCK cells at the end of the experiment (dashed line in A) divided by the length of a straight line from the top to the bottom of the collision (solid line in A). Data represent mean measurements of three experiments. Student’s t test was used to analyse the data, **p<0.005. **D** Cell speed (μm per hour) at which RasV12 cells treated with PBS (black) or Wnt5a (yellow) and DMSO or WNT-974 migrate. The data represent mean +/- s.d. from three independent experiment. Student’s t test was used to analyse the data. *p=0.0156; **p=0.0058. **E** Mean fluorescence intensity of CD44 at the boundary between RFP positive and RFP negative cells in vehicle or WNT-974 treated KrasG12D (KC) tissues harvested at the end of the treatment (see figure 5F). CD44 fluorescence was quantified and reported as in figure 4I. Control quantification is the same used in supplementary figure 4F. SuperPlot shows all the quantified ROIs (smaller circles) and the mean for each mouse (n=3 samples; larger circles - blue, green, and yellow). The graph shows mean and +/- s.d. for the three mice. Student’s t test was used to analyse the data. *p=0.0467. **F** Representative images of endogenous RFP fluorescence in PC pancreas tissue sections harvested 28-days post treatment with vehicle or WNT-974. Scale bar, 500µm. **G** RFP fluorescence per tissue area in PC mice treated with WNT-974 or vehicle and relative to vehicle treated tissues. Data represent mean and +/- s.d. per mouse. Student’s t test was used to analyse the data, ***p<0.0005. **H** Schematic summarising the main findings of the study. Illustration created with BioRender.com.

